# Dynamical Fingerprints of Rare-Event Transitions from Transition Corridors

**DOI:** 10.64898/2026.11.09.765370

**Authors:** Ali Sharifian, Alexander Schug

## Abstract

Rare transitions are central in science, yet their mechanisms are obscured by the high dimensionality of complex dynamics, a challenge particularly pronounced in biomolecules where rare conformational transitions underlie functional processes. A transition-conditioned dynamical fingerprint is introduced for a rare *A* – *B* event, in which the transition-relevant subset is targeted rather than a global decomposition and the problem is treated dynamically. Within a selected transition corridor, barrier-crossing motion is extracted and microscopic players are ranked through a corridor-conditioned informativeness measure, after which the highest-ranked players are subjected to intervention tests of their transition leverage using the REST2 scaling scheme, metadynamics, and targeted mutation. Across three benchmark proteins, targeting these players facilitates basin-to-basin transitions while maintaining high fidelity to the prescribed transition corridor, providing handles for mechanistic analysis, enhanced sampling, control, and design.

## Introduction

Rare events, despite their low probability, often determine the effective behavior of complex systems, from chemical kinetics to stochastic dynamics. In metastable regimes, weak fluctuations can induce transitions between long-lived states. ^1,2^ Because these transitions are typically inaccessible to direct simulation or experiment, their analysis requires specialized rare-event simulation and sampling methods. ^3–7^ Beyond characterization, a central objective is control: suppressing unwanted transitions or promoting desired ones.

Rare conformational transitions underlie biomolecular function, ^8^ including folding, ligand binding, allostery, and catalysis, because proteins populate ensembles of interconverting states. ^9–11^ Quantitative characterization of these transitions can enable strategies to modulate function. ^12^ Although modern simulations and experiments increasingly resolve such events, ^13–16^ mechanistic interpretation remains difficult because the relevant signal is embedded in high-dimensional dynamics. Progress therefore requires informative state representations, pathway and kinetic analyses, and identification of microscopic determinants with causal influence on the transition.

A substantial literature therefore targets low-dimensional representations (collective variable (CV)) of conformational dynamics. Despite their different criteria, the broader objective is the same: to obtain a compact representation in which relevant metastable states can be resolved. Early linear methods such as principal component analysis (PCA) identify dominant structural fluctuations, ^17^ but variance is not a reliable surrogate for slow kinetics. Time-correlation-based approaches, most notably time-lagged independent component analysis (tICA), instead target slow dynamical modes and underpin kinetic model construction. ^18,19^ Nonlinear manifold-learning and deep-learning methods extend such representations to more complex conformational landscapes. ^20–24^ Beyond representation learning, data-driven optimization approaches can also adapt biomolecular simulation parameters and protocols to targeted objectives. ^25^

Complementary to methods that seek compact representations of the conformational ensemble, rare-event and kinetic frameworks focus the description on the transitions themselves: how long-lived states are connected, which pathways carry reactive motion, and at what rates. Here, reduced descriptions serve a different purpose: states and network discretizations, or interfaces, milestones, and progress variables, are used to organize reactive dynamics and resolve transition mechanisms and kinetics. When these objects are defined in a chosen feature space, the resulting description can therefore depend on that representation. Transition Path Theory (TPT) provides a rigorous framework for characterizing reactive dynamics between prescribed sets in terms of committors, reactive fluxes, transition rates, and dominant pathways, and is commonly applied to Markovian models inferred from simulation data, including Markov state models (MSMs). ^26–28^ Related approaches characterize rare transitions through path ensembles, interfaces, or first-passage statistics, including TPS, ^29^ TIS, ^30^ FFS, ^31^ and milestoning. ^32^ Several methods further connect reduced-coordinate construction with rare-event sampling by optimizing or learning coordinates specifically for the transitions of interest. ^33,34^

Other approaches aim to identify microscopic players using residue-resolved, directly interpretable signals. Information-theoretic methods quantify statistical dependence among degrees of freedom, for example through mutual-information residue networks and multivariate extensions, to highlight coupled residues and putative communication pathways. ^35,36^ Other methods cast player identification as model reduction, selecting coordinate subsets that maximize information retention under KL-based objectives and thereby producing per-residue information fields. ^37,38^ In parallel, dynamical-network approaches construct residue interaction graphs from contacts or dynamical couplings and identify hubs and bridges through centrality or community structure. ^39^

Thus, these approaches can remain ensemble-pooled descriptions: even when slow dynamics, probability flow, or dynamical/information signals are incorporated, globally aggregated statistics may primarily preserve state separation, kinetic, or network structure, while the transition-local motion and informative players of a designated A–B barrier-crossing event need not be isolated; moreover, complex folding landscapes can support multiple competing pathways. ^40^ Beyond this, causal leverage over that event is not established by these criteria alone.

Also, recent deep-learning advances have enabled highly accurate prediction of near-native protein structures for many proteins, ^41,42^ yet robust generation and independent validation of biologically relevant alternative conformations—and, crucially, the mechanisms and kinetics connecting them—remain limited. AlphaFold-like models can sometimes be steered toward alternative states, ^43–45^ but enumerating states alone does not establish whether interconversion is kinetically accessible, which pathway is taken, or which microscopic elements control barrier crossing. This limitation is consequential for protein engineering and functional design: a desired metastable state may be predicted, while the corresponding transition may proceed by an unintended mechanism or remain too slow on experimental timescales. These limitations sharpen the need for transition-aware analyses that move beyond static structure prediction by resolving informative representations and microscopic determinants of functional dynamics.

Here, a transition-conditioned viewpoint yields a dynamical fingerprint of a designated *A* – *B* rare event by isolating the barrier-crossing signal—typically sparse and easily obscured—from equilibrium fluctuations and unrelated interconversions that confound global analyses. The fingerprint provides two actionable outputs: a compact representation of the collective barrier-crossing rearrangement and an event-specific ranking of microscopic players that are both informative in the transition-conditioned representation and endowed with intervention-tested causal leverage. Conceptually, this provides an operational analog of Maxwell’s demon, whereby information extracted from the ensemble guides spatially selective actuation to facilitate the desired rare event. These outputs furnish compact coordinates for analysis or biasing, identify candidate residues for mutagenesis or site-specific labeling, and provide a principled route to targeted coarse-graining and control. This perspective is broadly transferable to high-dimensional rare-event dynamics beyond proteins.

## Theory

A transition-conditioned approach is introduced for a specified rare event *A* – *B*. Global collective variables are first chosen to distinguish metastable basins and define a free-energy surface. A connecting path and a finite corridor around it are then identified, thereby selecting the configurations most relevant to the transition event. Dimensional reduction conditioned on this corridor is subsequently performed to identify the degrees of freedom most informative for barrier crossing, namely the dominant transition-local displacements and the associated slow coordinates. The resulting dynamical fingerprint comprises a low-dimensional representation of the transition motion, a ranking of microscopic players by their corridor-conditioned informativeness, and an intervention-based assessment of transition leverage.

### Global collective variables for representing metastable states

The trajectory is first represented in a CV space that separates and characterizes the metastable states relevant to the transition. In the present work, three families of global CVs are considered: RMSD/*R*_*g*_, PCA-based CVs, and tICA-based CVs. These choices are motivated by computational efficiency, empirical robustness, and broad applicability across diverse proteins. For visualization and free-energy estimation, each family is reduced to a two-component representation from which the landscape of basins and barriers is obtained. Detailed definitions and implementation are provided in Supporting Information 1, Sec. S1.1.

For each frame *t*, the resulting two-component representation is denoted 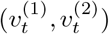. A kernel density 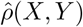 is then estimated on a grid, and the corresponding free-energy surface, in units of *k*_*B*_*T*, is defined as

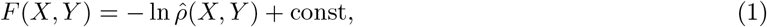

where the additive constant is chosen such that min *F* = 0.

### From free-energy landscapes to transition tubes

A transition corridor is defined in the selected two-dimensional CV free-energy landscape. After discretizing the landscape, a minimum-barrier path connecting the user-specified metastable basins is identified, motivated by *P* (*s*) *∝* exp[*− βF* (*s*)], so that lower-free-energy regions carry larger equilibrium weight and lower-barrier routes provide a natural approximation to statistically favorable transition channels in the projected landscape. This representation is computationally tractable, readily interpretable, and directly usable for subsequent mechanistic analysis; other criteria could define different paths and change the statistical or reactive relevance of the resulting route, while leaving the remaining method unchanged.

In the present work, the two-dimensional free-energy surface is represented on a rectangular grid *{*(*X*_*i*_, *Y*_*j*_)*}* with free-energy values *F*_*ij*_. The user-specified start and end collective-variable coordinates, corresponding to the metastable states of interest, are mapped to the nearest grid nodes. A minimum-barrier path *P* = *{p*_1_, …, *p*_*L*_*}* is then obtained on the grid, where *L* =|*P*| is the number of path nodes. Let 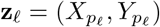, for *l* = 1, …, *L*, denote the corresponding path coordinates. A finite corridor is defined around this path through geometric and free-energy tolerances, such that only configurations sufficiently close to the path and not excessively high in free energy are retained. Simulation frames are mapped to their nearest grid nodes in collective-variable space and retained only if those nodes lie within the corridor. The retained frames are then assigned to their nearest path positions, thereby defining the transition tube used in the subsequent analysis. In this construction, the path and corridor are defined directly from the free-energy landscape, yielding a direction-free progress coordinate along the transition tube. Details are provided in Supporting Information 1, Sec. S1.2.

### Corridor-conditioned dimensional reduction for motions and players

A corridor-conditioned analysis is performed to identify both the collective coordinates that compactly represent the designated transition motion, referred to as Motion, and the microscopic structural elements, referred to as Players, that are most informative for it. The underlying intuition is that relevant players should display large variation within the transition-path corridor while also changing in a temporally coherent manner rather than as rapidly fluctuating noise. In the mathematical formalism, this intuition is expressed through an informativeness criterion that combines variance, measuring the magnitude of structural change, with slowness, measuring its persistence along the transition. Player rankings are then defined by this corridor-conditioned informativeness and are complemented by a separate intervention-based assessment that evaluates which highly informative players exhibit actionable leverage on the transition under controlled settings.

#### Corridor-conditioned Informativeness

To quantify how strongly a subset of coordinates *X*_*S*_ predicts transition progress on the corridor ensemble 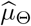, informativeness is defined as the explained-variance trace functional (full derivation and general form in Supporting Information 2),

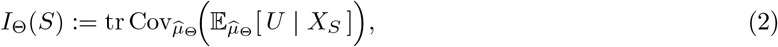

where *U* = (*U* ^(1)^, …, *U* ^(*r*)^) *∈* ℝ^*r*^ denotes the retained slow progress coordinates. A short derivation is obtained by adopting the best affine-predictor formulation for the conditional mean,

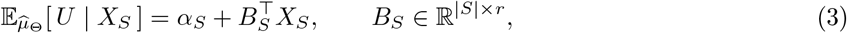

where *B*_*S*_ denotes the best affine predictor under 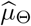.

With 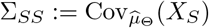, the constant term in the affine predictor does not contribute to covariance, and therefore

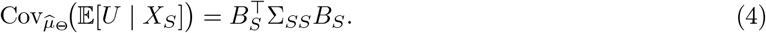

The optimal affine predictor satisfies the normal equations

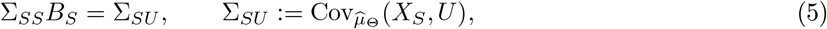

so that 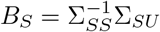 when Σ_*SS*_ is invertible. Substituting into Eq. (4) yields the set-level form

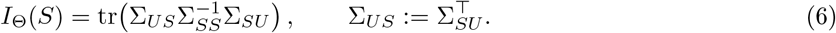

In a decorrelated ranking basis, Σ_*SS*_ is diagonal and the trace becomes additive,

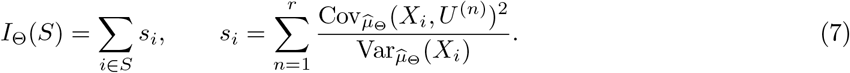

Specializing to PCA-score coordinates *Y* with

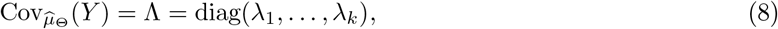

and writing the retained slow coordinates in that basis as the tICA components

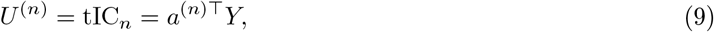

Substitution into the additive score gives

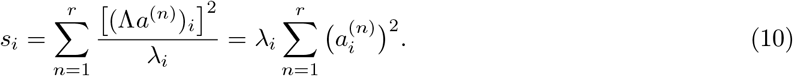

i.e. PCA variance times the sum of squared tICA loadings.

#### Players: Informative structural elements

Microscopic players are identified as structural elements with large corridor-local fluctuations and coherent contributions to the retained slow transition coordinates. Let *N*_*a*_ denote the number of retained atoms, and let 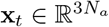 be the rigid-body-aligned Cartesian coordinates at frame *t*. For the corridor-frame set *T*_path_, with *T*_path_ = |*T*_path_|, the corridor mean is 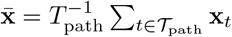. The centered corridor data matrix 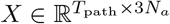 is assembled from 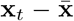, and its covariance is diagonalized as *C* = (*T*_path_*−* 1)^*−*1^*X*^*⊤*^*X* = *V* Λ*V* ^*⊤*^, with Λ = diag 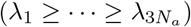.

The top-*k* loadings Φ = *V*_(:,1:*k*)_ and Λ_*k*_ = diag(*λ*_1_, …, *λ*_*k*_) are retained. The retained dimension was chosen to be large enough to preserve the transition-tube morphology upon reconstruction, but small enough to avoid dilution of the back-projected representation by broader non-transition fluctuations. In this work, 40% of the available modes were retained; data-adaptive truncation will be considered in future work.

For time-lagged analysis, each trajectory frame is first represented in this corridor-derived subspace as

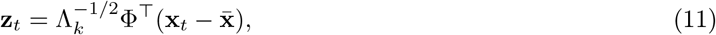

and the resulting full time-ordered score series is mean-centered before tICA, giving 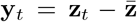 with 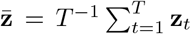.Thus, tICA is applied to the full trajectory only after projection onto displacement modes learned from the selected transition corridor. The instantaneous and time-lagged covariance matrices are estimated from the centered pairs (**y**_*t*_, **y**_*t*_) and (**y**_*t*_, **y**_*t*+*l*_), respectively, and the retained slow modes *W* = [**w**_1_, …, **w**_*r*_] are obtained from

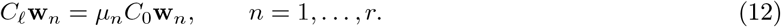

For Cartesian interpretation, the retained tICA modes are represented as 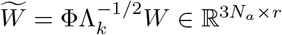. In the score basis 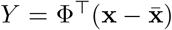, with the same mean-centering implicit, the retained slow coordinates are equivalently written as tIC_*n*_ = *a*^(*n*)*⊤*^*Y*, with 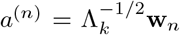. The corridor-conditioned informativeness derivation then gives 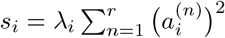, so that transition-local variance is weighted by coherent contribution to the retained slow transition coordinates. After Cartesian interpretation of the retained representation, the component scores associated with atom *α* are denoted *s*_*α,x*_, *s*_*α,y*_, and *s*_*α,z*_. The atom-level player score is 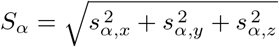, and atoms with larger *S*_*α*_ are ranked as more informative microscopic players for the selected transition.

#### Motions: Representation of the Designated Transition

Motion collective coordinates are constructed to provide a compact representation of the designated transition. The motion representation uses the collective contribution of all retained atoms through the orthonormal basis of Cartesian displacement modes obtained from the covariance of corridor frames. The full time-ordered trajectory is then represented in this fixed corridor-derived basis, and slow collective coordinates are extracted from the resulting score time series to describe progress along the transition tube.

Let 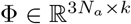 denote the *k*-dimensional orthonormal displacement basis obtained from the corridor-frame covariance analysis, and let Λ_*k*_ = diag(*λ*_1_, …, *λ*_*k*_) denote the corresponding retained eigenvalue matrix. Each trajectory frame is represented by the corridor-fitted score 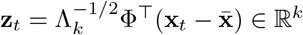. The projected full trajectory is then mean-centered before time-lagged analysis, giving 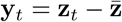, with 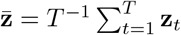.

The slow coordinates are obtained in this corridor-derived coordinate system and are used here as collective progress coordinates. With **w**_*n*_ denoting the retained tICA modes, the unscaled slow coordinates are tIC_*n*_ 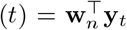, with *n* = 1, …, *r≤ k*. For the final motion representation, each slow coordinate is weighted by its corresponding tICA eigenvalue, WtIC_*n*_(*t*) = *µ*_*n*_tIC_*n*_(*t*), with *n* = 1, …, *r*. The *r*-dimensional representation of the designated transition is defined by these eigenvalue-weighted slow coordinates. The leading coordinates therefore provide a low-dimensional description of transition progress within the corridor-derived Cartesian displacement subspace.

#### From corridor informativeness to intervention-supported transition leverage

Corridor-conditioned informativeness was used to nominate players predictive of progress within the selected transition tube, but ranking alone was not interpreted as causality because predictive associations can reflect correlations or hidden confounding. ^46,47^ Since transition control is dynamical, causal support was assessed through robustness and controlled intervention rather than state-wise uncertainty alone, consistent with feedback-control and directed-information viewpoints. ^48–53^ This distinction was also motivated by our earlier work on histidine-kinase activation, from the characterization of inactive and active conformations ^54^ to the transition-specific dynamical analysis of WalK. ^55^ For WalK, an RMSD-based bias applied only to the

DHp domain produced an activation rearrangement comparable to that obtained when both the DHp and CA domains were biased, with a concurrent CA rotation of *∼*50^*°*^, only *∼*7^*°*^ smaller than in the both-domain-biased protocol. ^55^ This shows that localized elements can exert transition-specific dynamical leverage over a collective rearrangement. In the present framework, corridor-conditioned informativeness is therefore treated as a hypothesis generator that nominates candidates, whereas causal support is assessed separately through robustness across distinct corridor constructions and a dedicated controlled setting. Candidate players were therefore evaluated using two criteria for intervention-tested transition leverage: (i) robustness of the ranking across distinct CV-defined corridors, and (ii) controlled separation of strongly correlated candidates, here |*ρ*| *>* 0.7, with one representative used at a time and correlated alternatives substituted in separate simulations. Additional details are provided in Supporting Information 1, Sec. S1.3.

## Results and Discussion

Deep-learning structure predictors have transformed access to near-native stable states and can sometimes be steered toward alternative conformations. ^41,43–45,56^ The next bottleneck is mechanistic: without transition-specific dynamics, it is difficult to determine which alternatives are functionally relevant and how exchange occurs. ^57^ Here the focus is shifted from “which structures?” to “which transitions, motions, and players?”: a path-conditioned dimensionality-reduction approach extracts a low-dimensional barrier-crossing motion, ranks microscopic players by corridor-conditioned informativeness, and assesses transition leverage separately through controlled interventions.

### Theory-to-intervention tests of transition leverage

To operationalize the shift from structures to transitions, high-dimensional conformational dynamics of proteins are represented in a low-dimensional collective-variable free-energy landscape, where a finite-width corridor following the low-free-energy transition region defines the transition-conditioned ensemble. Analysis within this ensemble extracts the barrier-crossing motion, ranks informative atoms, and nominates intervention targets. Enhanced sampling restricted to a Hot-region of interest (ROI) of top-ranked atoms is expected to facilitate the transition while suppressing nonspecific off-corridor exploration, because these atoms are selected from low-free-energy transition ensemble and contribute coherently to motion along it. Accordingly, the corridor is treated as a prescribed transition region constructed around the selected low-free-energy path, rather than as a reactive pathway inferred from reactive-trajectory statistics. Players and interventions are instead evaluated by whether they promote *A*–*B* crossing while remaining within this region.

For each protein, free-energy surfaces were constructed in three global CV representations: RMSD/*R*_*g*_, PCA, and tICA (Figure 1). In each representation, the minimum-barrier path and corridor tube were identified using the Theory protocol, and the associated corridor frames were used for downstream motion and player analysis. The tICA-based landscape is shown in the main text for visualization, while the RMSD/*R*_*g*_ and PCA analyses are provided in Supporting Information 3, Sec. S3.2. The same corridor-conditioned analysis yields the motion representation in Figure 1(d): weighted tICs (WtICs), defined as eigenvalue-weighted slow coordinates obtained after projection into the corridor-derived Cartesian displacement basis. Players were ranked within each CV-defined corridor by corridor-conditioned informativeness. For each protein, the Hot-ROI was defined as the mutual high-rank atom set reproduced across the three CV constructions, providing robustness to the choice of global landscape used to define the tube (Supporting Information 1).

**Figure 1.**
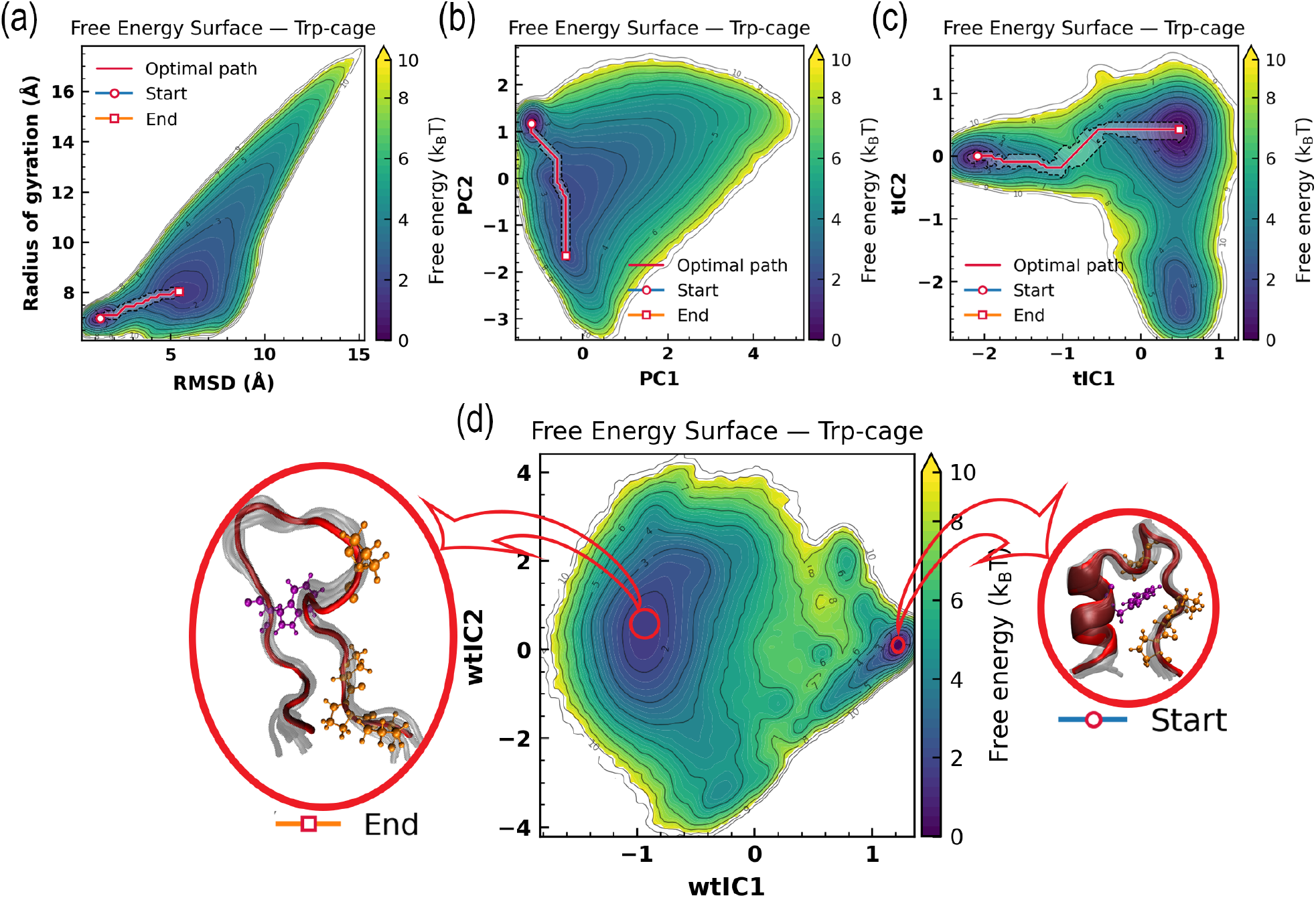
Free-energy landscapes of Trp-cage from a 208 µs simulation in three global CV representations: (a) RMSD–*R*_*g*_, (b) PCA, and (c) tICA, with the minimum-barrier path and transition corridor indicated. (d) Motion collective coordinates. The two metastable basins are defined by the start-cluster and end-cluster structures. For clarity, PRO is shown in orange and TRP in purple.

Transition leverage was tested by two complementary interventions. The REST2 scaling scheme was used to apply localized Hamiltonian scaling to the selected ROI, whereas multi-walker well-tempered metadynamics was used as a CV-based sufficiency test by applying the same biasing protocol while varying only the biased atom set: all-atoms, Hot-ROI, or Random-ROI. ^58–61^ Transition facilitation and corridor fidelity were assessed using prespecified analyses. Validation used matched ROI definitions for the REST2 scaling scheme analyses: Hot-ROI was defined as the top-*N* atoms ranked by *S*_*α*_, Random-ROI as randomly selected *N* atoms, and Tail-ROI as the next *N* atoms after removing the top *M* . Additional controls included Path-PCA-ROI, Path-tICA-ROI, and G-tICA-ROI, defined from path-conditioned PCA, path-conditioned tICA, and global tICA rankings, respectively, together with Data-fraction tests in which *S*_*α*_ was recomputed from reduced reference data (Table 1). Matched REST2 scaling-scheme cohorts used identical schedules and simulation time budgets; outcomes are reported in Figures 2, 3 and 5. All tests were performed on Trp-cage, ^62,63^ BBA, ^64^ and villin headpiece; ^65^ BBA and villin results are summarized in Appendix, Sec. A.1. Random-ROI controls were evaluated using five independent seeds, whereas all other ROI selections were evaluated using three independent seeds. One realization per condition is shown for visualization only, while all quantitative comparisons are based on the independent replicate runs; the corresponding seed-resolved results are reported in Supporting Information 3. Long unbiased trajectories used to construct the corridors were taken from Anton datasets from D. E. Shaw Research. ^14^

**Table 1:**
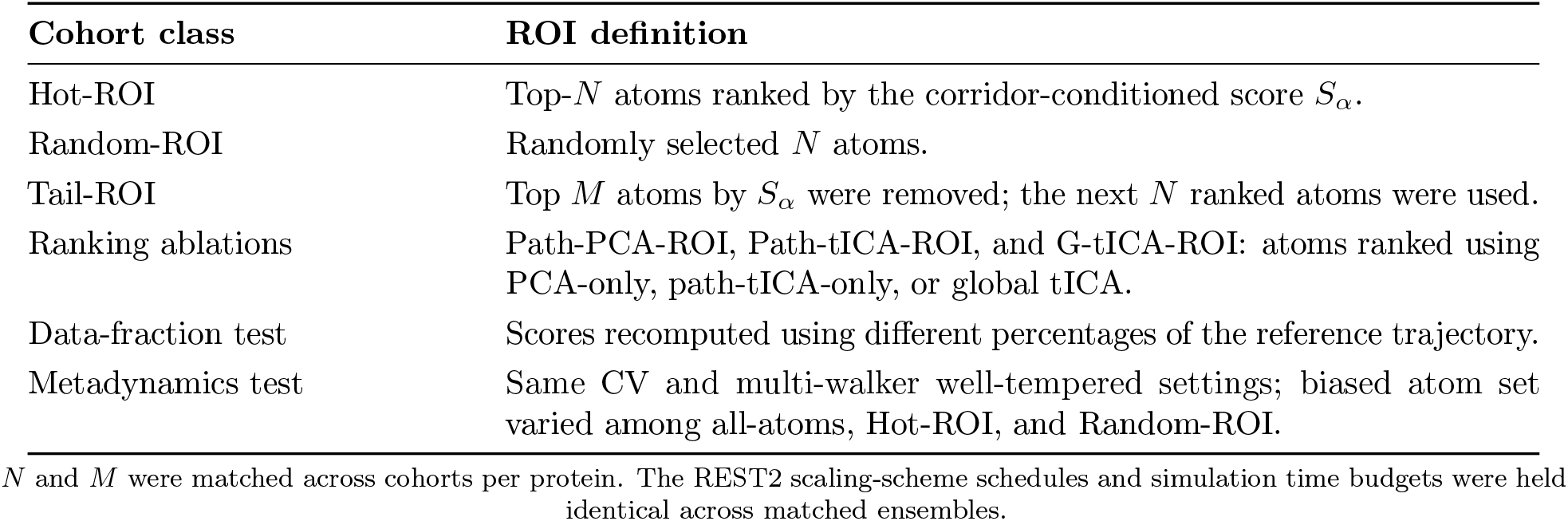
ROI cohorts used for validation.

**Figure 2.**
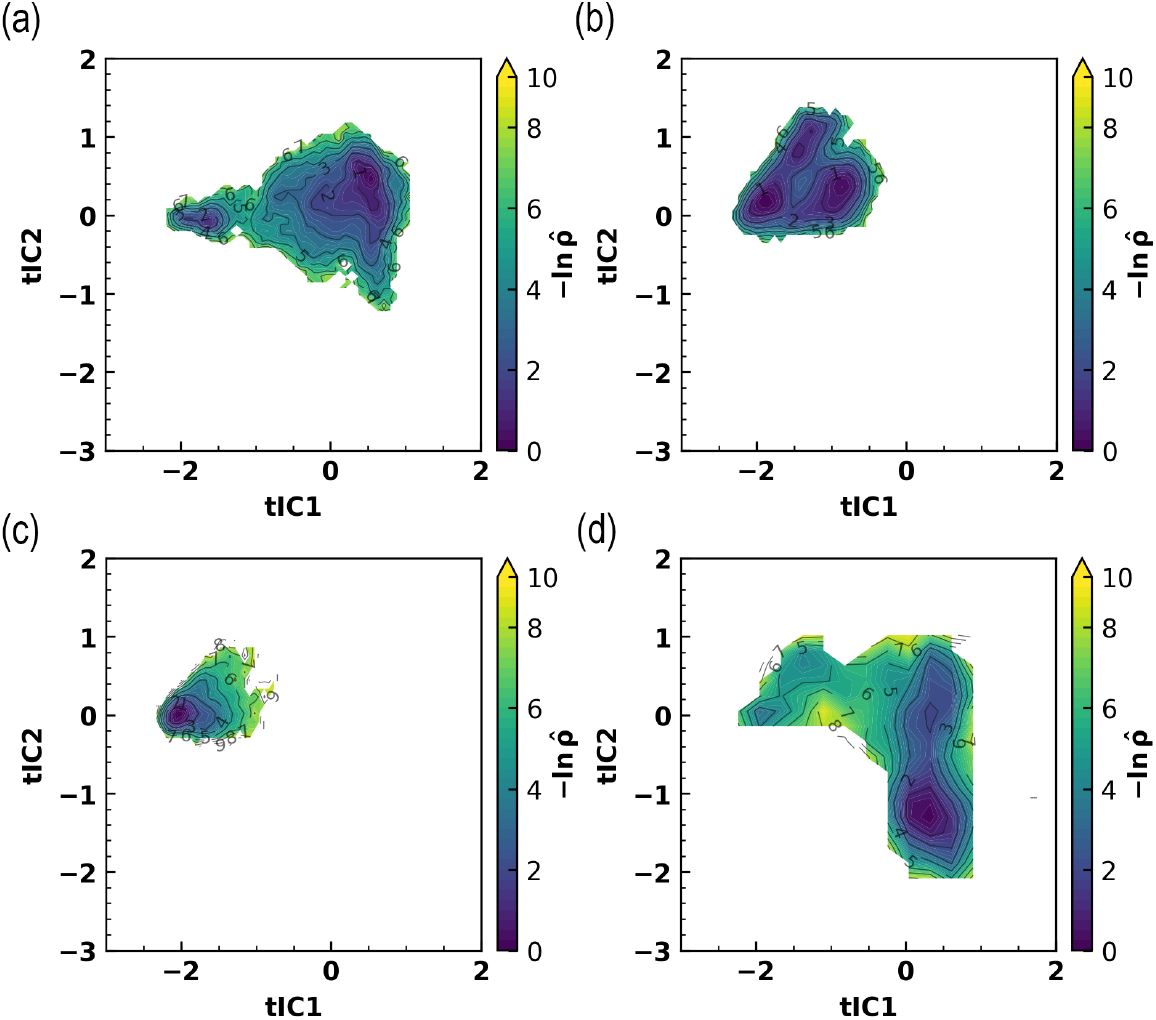
tICA-based sampling-density maps of Trp-cage computed from 100 ns simulations for (a) Hot-ROI, (b) Random-ROI, (c) Path-PCA-ROI, and (d) Path-tICA-ROI.

**Figure 3.**
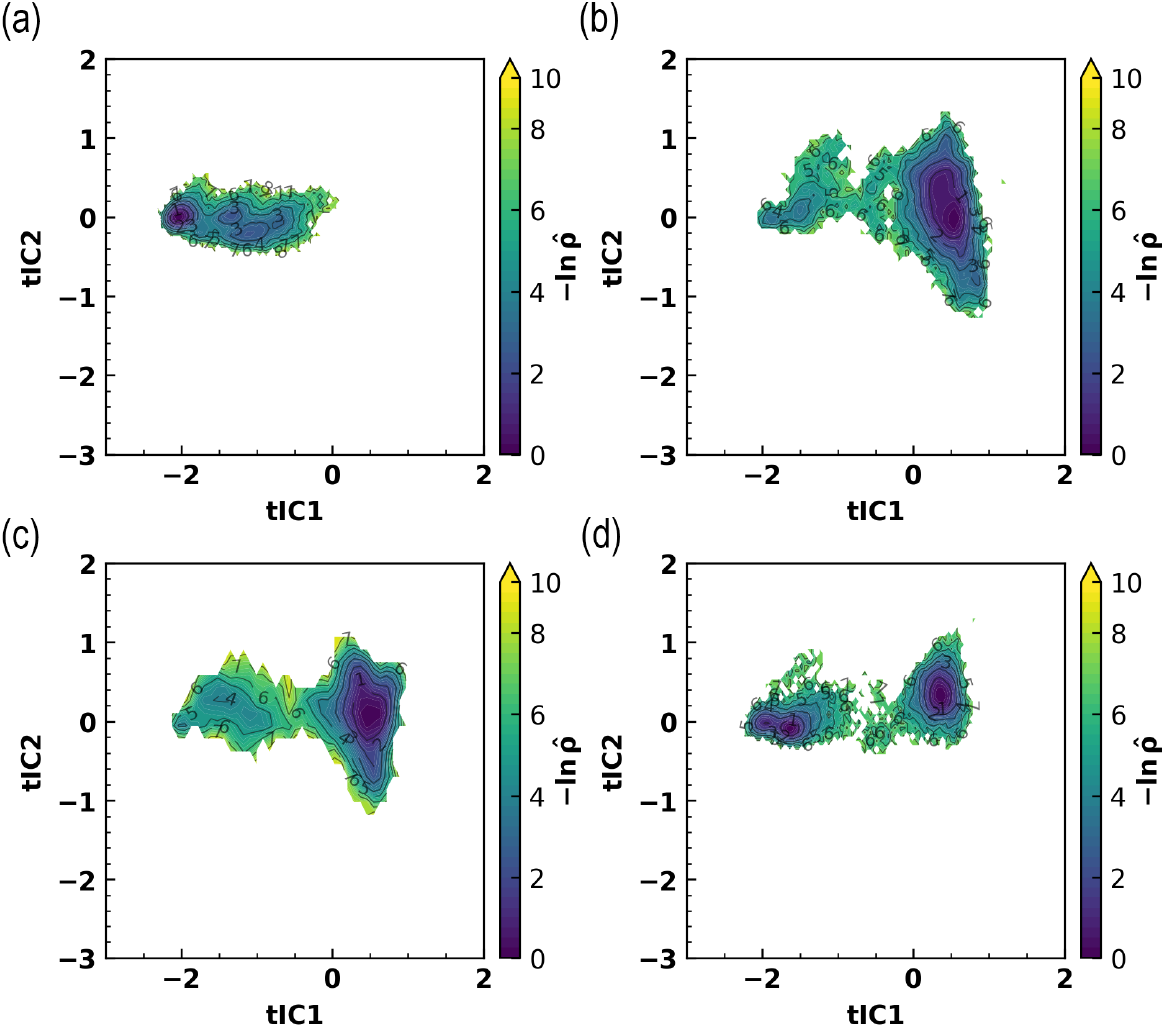
tICA-based sampling-density maps of Trp-cage computed from 100 ns simulations for (a) 4% Hot-ROI, (b) G-tICA-ROI, (c) Tail-ROI, and (d) 20% data-fraction.

**Figure 4.**
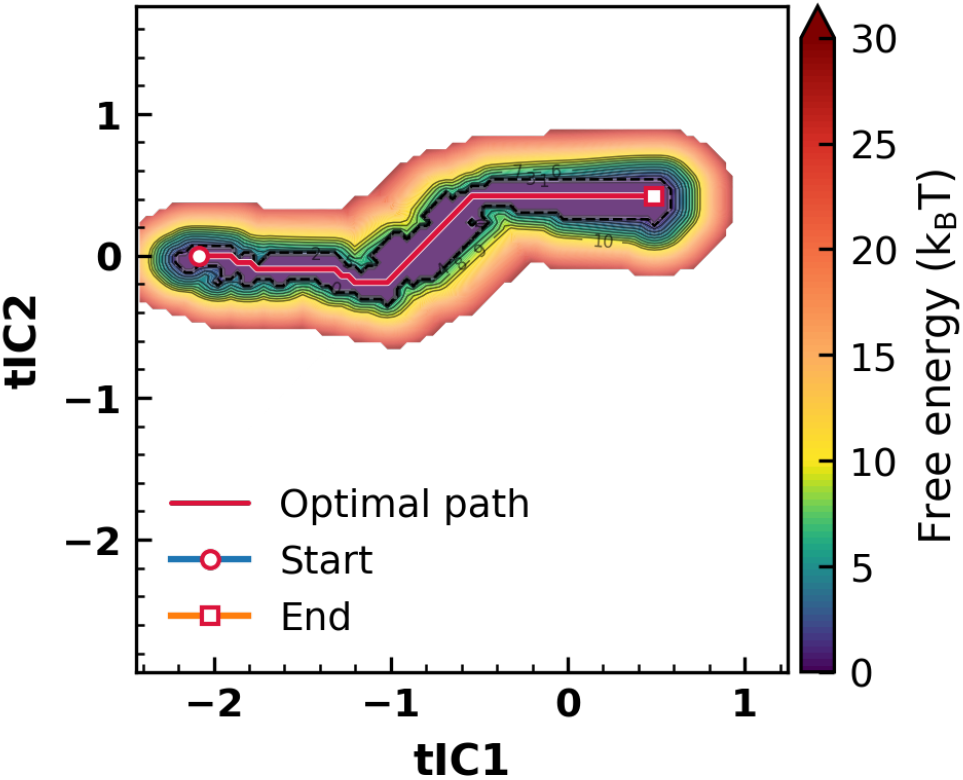
Corridor-weighted reference map, showing the prescribed transition corridor and the increasing off-corridor penalty used in the fidelity analysis.

**Figure 5.**
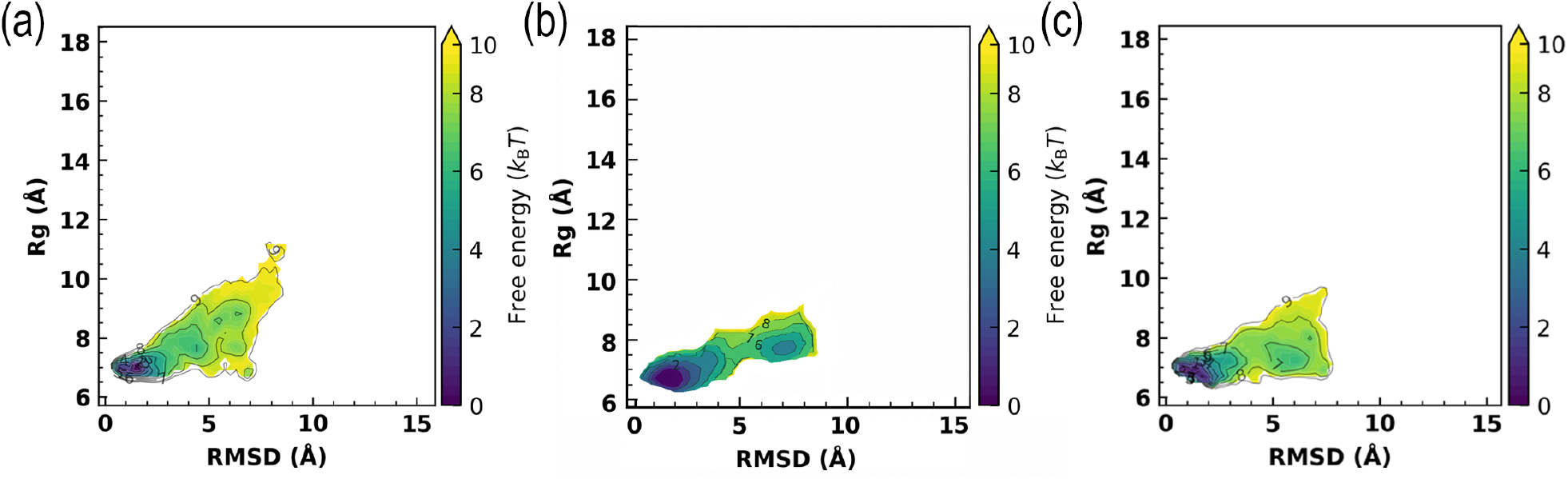
RMSD–*R*_*g*_ free-energy profiles from metadynamics under three biasing choices: (a) all atoms, (b) Hot-ROI, and (c) Random-ROI.

All systems were prepared and simulated in GROMACS 2019.6 ^66,67^ patched with PLUMED 2.7.3, ^68,69^ using the CHARMM22* force field ^14,70,71^ with the TIP3P water model. ^72^ Nonbonded interactions used 1.2 nm cutoffs and PME electrostatics. ^73^ All equilibration and production simulations employed a 2 fs leap-frog integrator with LINCS constraints; ^74^ temperature was controlled with a velocity-rescaling thermostat, ^75^ and pressure, where applicable, with a Parrinello–Rahman barostat. ^76^ Systems were first energy-minimized and then equilibrated under the unscaled Hamiltonian by an 800 ps NVT simulation at *T*_ref_ = 330 K followed by an 800 ps NPT simulation, and then an additional 800 ps relaxation stage. The Hamiltonian scaling scheme of REST2 ^77^ was applied to the selected ROI with *λ* = 0.4238 and run as a single trajectory without replicas or exchange attempts. Free energies from well-tempered metadynamics were obtained by reweighting the accumulated bias.

### Hot-ROI targeting outperforms baseline ROI strategies

Hot-ROI targeting was compared with Random-ROI, Path-PCA-ROI, and Path-tICA-ROI controls on Trp-cage, BBA, and villin headpiece using matched 100 ns simulations employing the REST2 scaling scheme. The resulting ensembles were projected onto the tICA landscape used for corridor construction, and the corresponding sampling-density maps are shown in Figures 2 and A-2.

Across all three proteins, Hot-ROI produced the most transition-competent and corridor-faithful sampling, reaching both metastable basins while enhancing sampling along the transition tube. In contrast, Random-ROI and Path-PCA-ROI often failed to reach the second basin, whereas Path-tICA-ROI could induce exchange but with increased off-corridor exploration; in villin, Path-PCA-ROI reached the second basin but with poorer corridor fidelity than Hot-ROI. These results support that combining corridor-local structural amplitude with slow dynamical coherence identifies players with transition-specific leverage.

Time-resolved trajectories in the same reduced representation are provided in Supporting Information 3, Sec. S3.3, Figures S3-6 and S3-7, showing the temporal progression of each intervention through the corridor and between metastable basins.

### Hotspot size, data fraction, and global tICA

Sensitivity of Hot-ROI targeting was tested under the same 100 ns REST2 scaling protocol and corridor/sampling-density diagnostics (Figure 3). Reducing the number of selected Hot-ROI atoms from the default 8% to 4% removed observable basin-to-basin transitions, indicating that sufficient hotspot size is required under fixed intervention strength; the hotspot fraction was therefore treated as a controlled hyperparameter. Tail-ROI, obtained by removing the top 4% ranked atoms and selecting the next 8%, and the Data-fraction test, in which scores were recomputed from 20% of the trajectory, both outperformed Random-ROI, Path-PCA-ROI, and Path-tICA-ROI controls, but showed lower corridor fidelity and higher apparent barriers than the full Hot-ROI. Separately, G-tICA-ROI produced broader exploration but reduced corridor specificity, consistent with global slow modes capturing fluctuations beyond the designated transition channel. These results indicate that the highest-ranked hotspots provide the strongest transition-specific leverage, while secondary, data-limited, or globally defined selections can remain effective but with diminished corridor control.

### Quantitative assessment of transition facilitation and corridor fidelity

Transition facilitation and corridor fidelity were quantified relative to a single corridor-weighted reference map. Conceptually, the aim of this analysis was not to reconstruct or compare the full free-energy landscape, but to examine the transition specifically through the prescribed corridor. Starting from the reference free-energy surface *F*_0_(*x, y*), a baseline map was therefore constructed so that the corridor itself remained the preferred region, while deviations away from it were increasingly penalized. The resulting corridor-weighted reference map is shown in Figure 4.

The baseline map was defined as

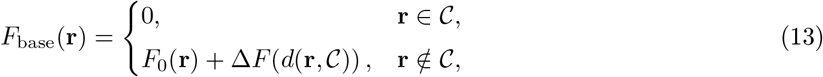

where *C* is the prescribed corridor and *d*(**r**, *C* ) is the Euclidean distance from grid point **r** to *C*. The off-corridor penalty was chosen as

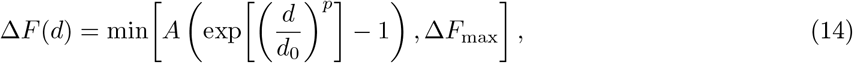

with *A* = 10, *d*_0_ = 0.3, *p* = 1, and Δ*F*_max_ = 120. This choice makes the corridor the preferred low-free-energy route and imposes a rapidly increasing cost away from it. Deviations from the baseline therefore reflect both loss of pathway fidelity and reduced occupancy along the corridor. Robustness to the corridor-weighting parameters was tested by varying *A* and *d*_0_ by 20%, both individually and jointly. The ranking of the ROI selections was unchanged, and the change in the score separation between the first- and second-ranked cases remained below 1.3%. The qualitative interpretation therefore remained robust to the choice of *A* and *d*_0_.

Each intervention was then compared with *F*_base_ using three complementary discrepancy measures. The Wasserstein (earth-mover) distance (EMD) ^78,79^ measures the minimum transport cost required to transform one probability distribution into another and therefore quantifies the overall spatial redistribution of probability mass. The two-dimensional Kolmogorov distance *K*_2D_ ^80^ instead quantifies the maximum absolute difference between the probabilities accumulated within corresponding quadrants of the two distributions, and therefore captures their strongest cumulative distributional discrepancy. The root-mean-square error (RMSE), in contrast, compares the two maps grid point by grid point and therefore quantifies pointwise differences in map values rather than the spatial redistribution of probability mass. These three measures consequently provide complementary information about agreement with the corridor-centered reference. In all cases, smaller values indicate closer agreement with the corresponding aspect of the reference. For the quantitative values reported in this subsection, uncertainties after *±*denote standard deviations over three independent seeds. The Random-ROI, 4% Hot-ROI, and Path-PCA-ROI cases were not assigned values, because they did not reach the second metastable basin on the analyzed timescale and therefore did not realize a corridor-relevant transition under the present criterion.

For simulations that sampled both metastable basins, the measured Wasserstein distances were

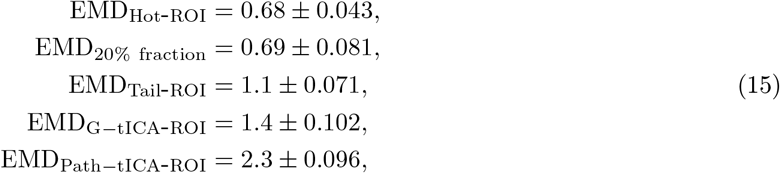

and the corresponding *K*_2D_ values were

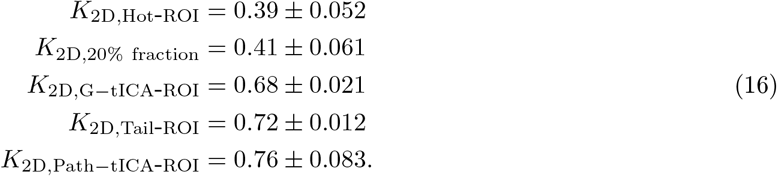

The corresponding pointwise differences in map values were quantified using the RMSE, yielding

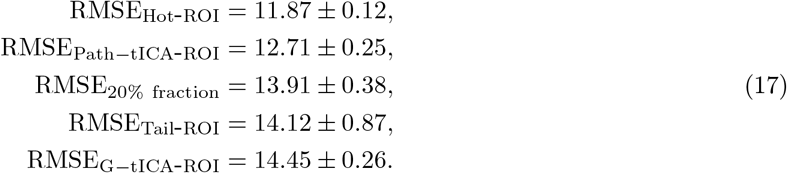

The three measures distinguish complementary aspects of corridor fidelity. Relative to the corridor-centered reference, the Hot-ROI yields the lowest value for all three metrics: the smallest overall redistribution of probability mass (EMD), the smallest maximum difference in accumulated quadrant probability (*K*_2D_), and the closest pointwise agreement in map values (RMSE). The agreement of all three criteria therefore identifies the Hot-ROI as the intervention that most faithfully reproduces the prescribed transition pattern.

The 20% data-fraction case further illustrates why the three measures should be considered together. Its Wasserstein and *K*_2D_ values are very close to those of the Hot-ROI and clearly lower than those obtained for the other alternative ROI selections. Comparatively, this indicates that the 20% data-fraction case reproduces the overall transition-associated probability distribution substantially better than the other alternatives and approaches the behavior obtained with the full Hot-ROI selection. Its larger RMSE, however, separates it more clearly from the Hot-ROI, indicating weaker pointwise agreement with the corridor-weighted reference. Thus, although the reduced-data selection captures the transition and its overall distributional organization comparatively well, it does not reproduce the detailed corridor-centered map as faithfully as the Hot-ROI. The stronger separation revealed by the RMSE, despite the close Wasserstein and *K*_2D_ values, therefore highlights the additional information provided by the pointwise assessment of corridor fidelity.

The Path-tICA-ROI shows a different balance between these measures. Its Wasserstein and *K*_2D_ values are substantially larger than those of the Hot-ROI and the 20% data-fraction case, consistent with its broader sampling and greater excursions away from the prescribed transition region. At the same time, it gives the second-lowest RMSE, indicating closer pointwise agreement with the corridor-weighted reference than the remaining ROI selections, though still clearly poorer than the Hot-ROI. The Path-tICA-ROI therefore remains comparatively closer to the corridor-centered reference in pointwise terms than several other alternatives, while its larger Wasserstein distance indicates a broader spatial redistribution of sampled density, consistent with more extensive low-occupancy exploration away from the corridor.

Taken together, these results show that distributional and pointwise agreement capture distinct but complementary aspects of corridor fidelity. Across all three metrics, the Hot-ROI is the only selection that consistently performs best, remaining closest to the prescribed transition tube in both distributional organization and pointwise map agreement. This supports the interpretation that the selected hotspots capture a transition-specific dynamical fingerprint rather than merely reproducing a generic low-dimensional transition pattern. Notably, the 20% data-fraction ranking still approaches the Hot-ROI in the distribution-based measures and outperforms the other alternative selections in this respect, indicating that a substantial part of the transition-specific signal can already be recovered from a markedly reduced amount of input data. Its larger RMSE, however, also shows that the full Hot-ROI selection retains a clear advantage in pointwise agreement with the corridor-centered reference.

### Hot-ROI biasing facilitates corridor-consistent transitions

Mechanistic sufficiency of the Hot-ROI was tested with well-tempered metadynamics by biasing the same RMSD–*R*_*g*_ coordinates while varying only the atom set used to define them: all backbone atoms, Hot-ROI, or Random-ROI (Figure 5). All-atom biasing produced broad, non-specific exploration, and Random-ROI biasing promoted slower exploration without consistent localization to the corridor. In contrast, Hot-ROI-restricted biasing drove efficient basin-to-basin transitions while remaining concentrated along the corridor-defined pathway. The same contrast was already visible at shorter accumulated simulation times (Supporting Information 3, Sec. S3.4, Figure S3-9). These results show that Hot-ROI-restricted CVs can facilitate the transition without converting the bias into generic conformational exploration, supporting the transition-specific leverage of the ranked Hot-ROI players.

### Transition-guided mutations perturb barrier crossing as predicted

As mutations can easily shift the balance between multiple conformations, ^81^ a mutation-level test was performed in Trp-cage by methylating ASP1, ASP9, and SER20, three highly ranked residues selected from the dynamical fingerprint. Methylation was used as a local steric perturbation to reduce conformational freedom at sites predicted to influence passage along the corridor, and the mutant system was evaluated using the same multi-walker well-tempered metadynamics settings as above (Figure 6). The mutant showed an increased inter-basin barrier relative to the long unbiased reference, and the density-difference map revealed depletion of the corridor-like connecting region together with enhanced sampling of flanking off-corridor regions. This redistribution indicates that the mutation makes the corridor less favorable and diverts sampling toward off-corridor routes, consistent with the interpretation that the highly ranked players carry dynamical leverage over barrier crossing.

**Figure 6.**
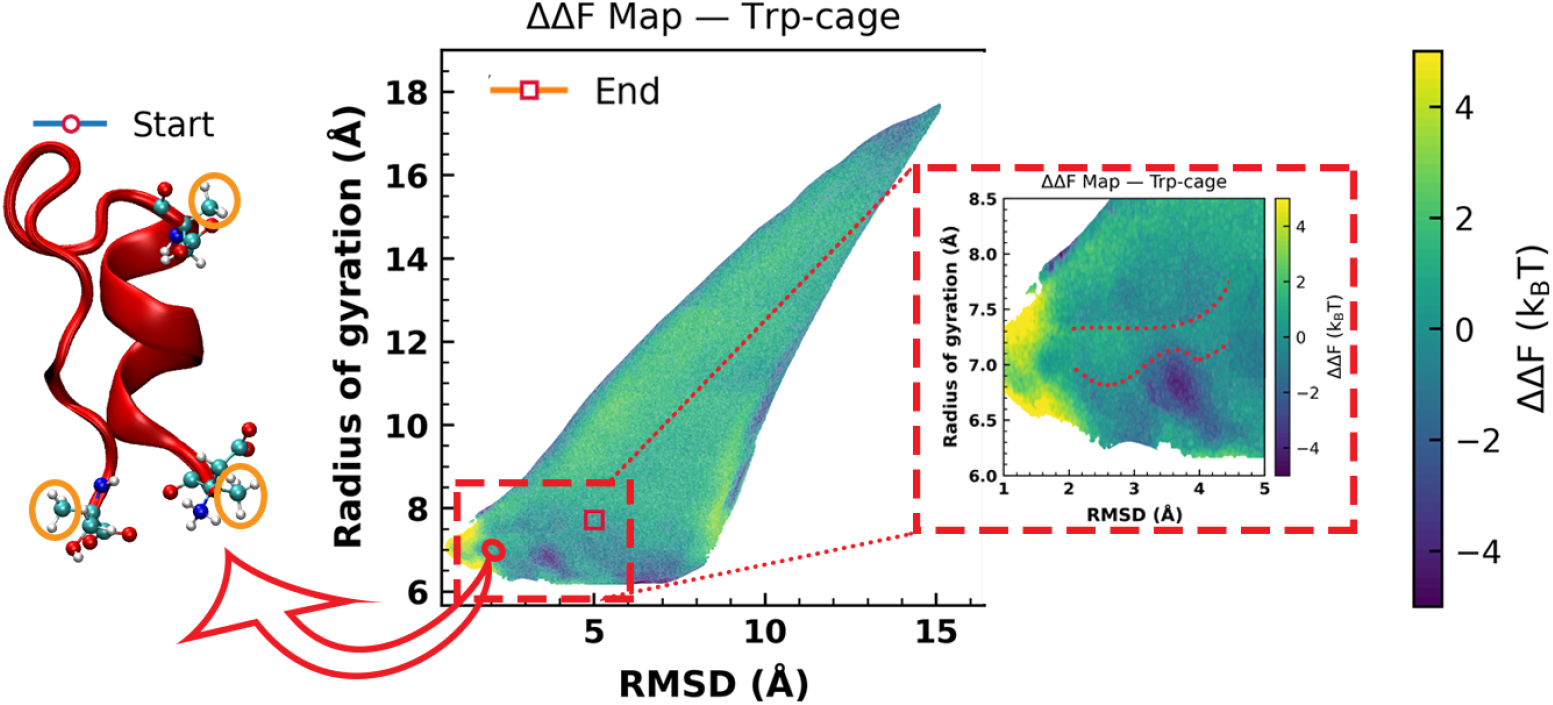
RMSD–*R*_*g*_ free-energy difference profiles for Trp-cage, defined as ΔΔ*F* (RMSD, *R*_*g*_) = Δ*F*_mutant_(RMSD, *R*_*g*_) *−* Δ*F*_ref_ (RMSD, *R*_*g*_), where Δ*F*_ref_ is the long unbiased reference free-energy surface. Positive ΔΔ*F* indicates regions that are less favorable in the mutant, whereas negative ΔΔ*F* indicates regions that are more favorable. The low ΔΔ*F* corridor shown in the red dashed box was obtained using the same corridor selection procedure described in the theory section. The three methylated residues are highlighted by the orange ellipse.

## Conclusion

Rare-event dynamics was resolved by conditioning the analysis on transition corridors, where a general informativeness formulation was used to link transition progress to microscopic coordinates within the selected region. Barrier-crossing motion, informative players, and dynamical drivers were then separated by corridor-conditioned dimensional reduction, Hot-ROI ranking, and controlled intervention tests, supporting a transition-local view of biomolecular mechanism, with potential applicability to broader complex dynamical systems.

## Notes

The authors declare no competing financial interest.

## Data and Code availability

The long unbiased reference trajectories analyzed in this study were obtained from D. E. Shaw Research and should be requested directly from D. E. Shaw Research ^14^ in accordance with their data-access conditions. Simulation data generated in this study that support the reported intervention analyses are available from the corresponding author upon reasonable request. The analysis code used to generate the reported results is openly available at https://anonymous.4open.science/r/dynamical-fingerprints-B3EB.

## Supporting Information

The Supporting Information contains implementation details; the mathematical formulation of corridor-conditioned informativeness; and additional figures together with details of the validation and intervention tests.

## Acknowledgments

The authors thank Tomáš Kubař and Julian Herold for helpful discussions. The authors acknowledge the NHR Center NHR@KIT for providing computational resources on the HoreKa supercomputer under project ID p0026542. Support from the DFG-funded Research Training Group RTG 2450 / GRK 2450, Tailored Scale-Bridging Approaches to Computational Nanoscience, at KIT is gratefully acknowledged.

During manuscript preparation, the authors used AI-based tools (Claude Opus and GPT Astra) to improve the clarity, grammar, and organization of the manuscript text. The authors reviewed and edited all AI-assisted revisions and take full responsibility for the content of this work.

## A Appendix

### A.1 Additional benchmark results for BBA and villin

**Figure A-1:**
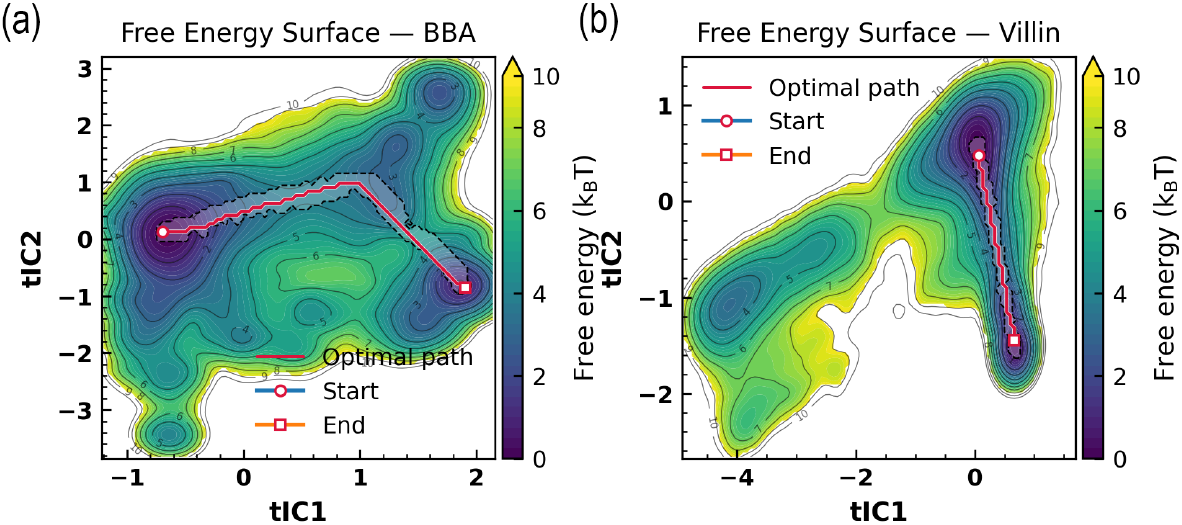
tICA-based free energy landscapes for (a) BBA (325 µs simulation) and (b) villin headpiece (125 µs simulation)

**Figure A-2:**
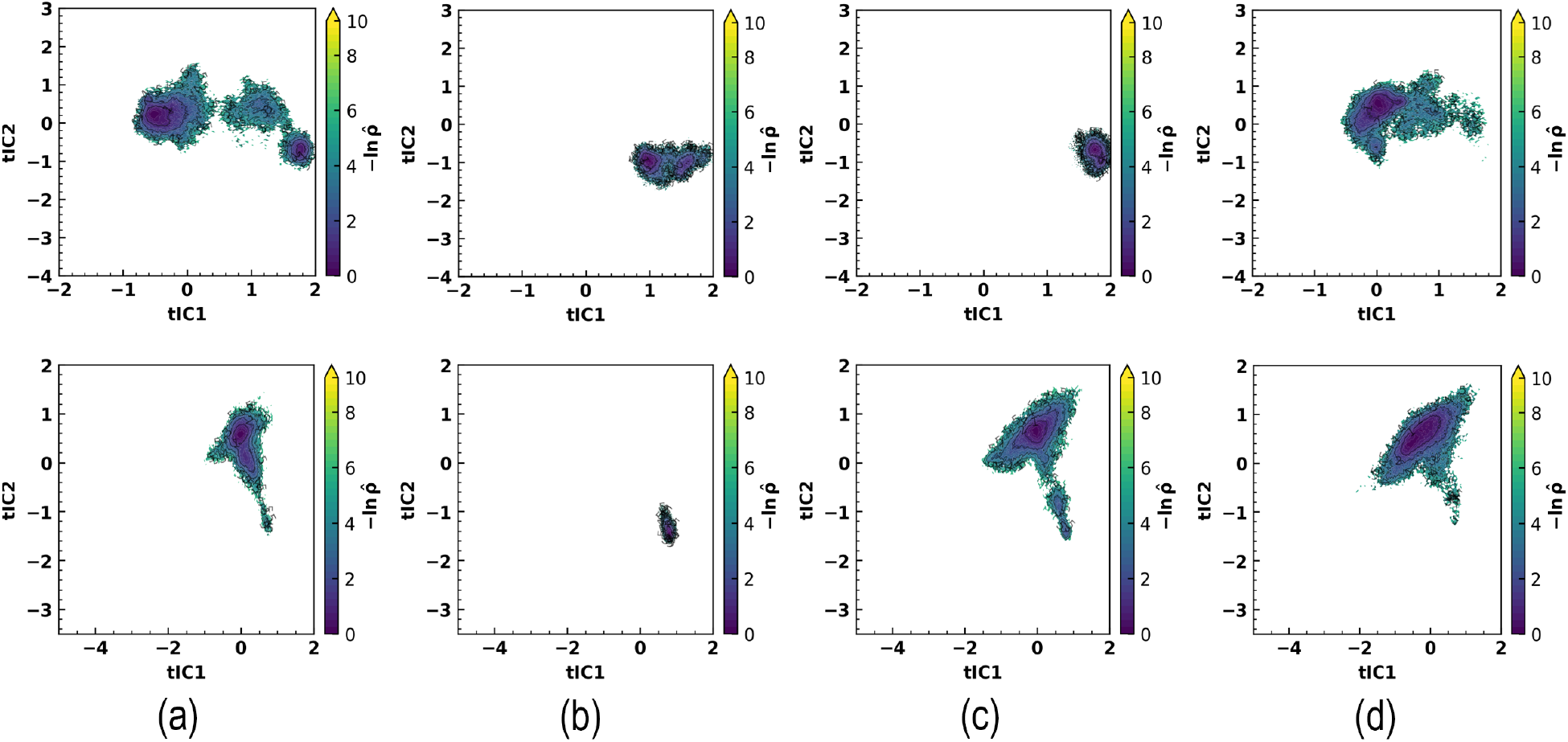
tICA-based sampling-density maps computed from 100 ns simulations for (a) Hot-ROI, (b) Random-ROI, (c) Path-PCA-ROI, and (d) Path-tICA-ROI. Rows correspond to BBA (top) and villin headpiece (bottom).

## Visual Abstract

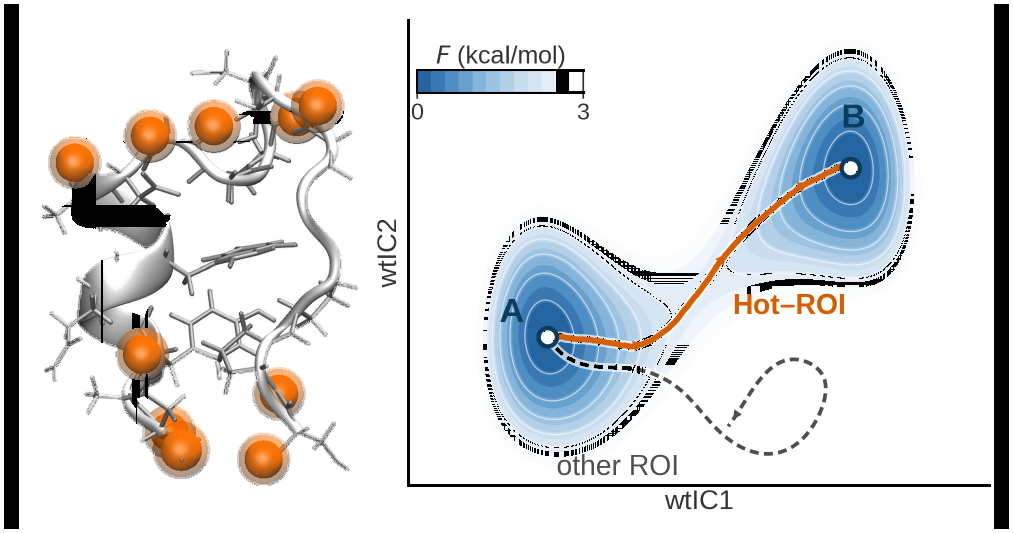

