## supplemental File for "Dynamical Fingerprints of Rare-Event Transitions from Transition Corridors"

**CV** collective variable

**FES** free-energy surface

**tICA** time-lagged independent component analysis

**TPT** transition path theory

**PCA** principal component analysis

**ROI** region of interest

### S1 Supporting Information 1

#### S1.1 Definitions and implementation of Global collective variables for representing metastable states

The trajectory is first mapped to a global collective variable (CV) space that separates the metastable states relevant to the transition. Three two-dimensional representations are considered: RMSD/$R_g$, PCA-based CVs, and tICA-based CVs, chosen for computational efficiency, robustness, and broad applicability. These representations are used to visualize the free-energy landscape and identify the basins and barriers.

Let $\{\mathbf{X}_t\}_{t=1}^{T}$ denote the Cartesian coordinates of the system, with $\mathbf{X}_t \in \mathbb{R}^{3N}$, and let $\{\tau_t\}$ denote the corresponding time stamps. In the present work, all global collective variables (CVs) are defined from $C_\alpha$-based structural descriptors, and each frame is represented by a two-component vector $(v_t^{(1)}, v_t^{(2)})$.

For visualization and free-energy estimation, the two-component representation is evaluated on a regular grid. A kernel density estimate $\hat{\rho}(X, Y)$ is constructed from the projected coordinates $(v_t^{(1)}, v_t^{(2)})$, and the corresponding free-energy surface is defined in units of $k_B T$ as

$$F(X, Y) = -\ln \hat{\rho}(X, Y) + \text{const}, \tag{S1-1}$$

where the additive constant is chosen such that $\min F = 0$.

Three CV families are considered.

(i) **RMSD/$R_g$ CVs.** A two-dimensional representation is defined by

$$v_t^{(1)} = \text{RMSD}(\mathbf{X}_t, \mathbf{X}_{\text{ref}}), \qquad v_t^{(2)} = R_g(\mathbf{X}_t),$$

where $\mathbf{X}_{\text{ref}}$ denotes the chosen reference structure. Here, RMSD is computed on the $C_\alpha$ atoms after optimal structural alignment, and $R_g$ denotes the corresponding radius of gyration.

(ii) **PCA-based CVs.** For each frame, the vector of $C_\alpha$ pairwise distances is constructed and standardized over the trajectory. Principal component analysis (PCA) is then performed on the resulting feature matrix, and the first two principal components are used as the two-dimensional CV representation,

$$(v_t^{(1)}, v_t^{(2)}) = (\text{PC1}_t, \text{PC2}_t).$$

(iii) **Global tICA-based CVs.** The same standardized $C_\alpha$ pairwise-distance features are used as input to time-lagged independent component analysis (tICA). A user-specified lag time is employed, and the first two global tICA components are taken as the CV representation,

$$(v_t^{(1)}, v_t^{(2)}) = (\text{tIC1}_t, \text{tIC2}_t).$$

If the lag is specified in frames, the corresponding physical lag time is obtained from the trajectory time step; conversely, if it is specified in physical units, the corresponding number of frames is used in the estimator.

These three CV families were selected because computational efficiency, empirical robustness, and broad applicability across diverse proteins were sought. Although only two-dimensional representations are used in the main text to define the free-energy surface and metastable-state geometry, higher-dimensional representations could also be considered.

**RMSD and radius of gyration ($C_\alpha$ definition)** RMSD was evaluated after rigid-body superposition to a $C_\alpha$ reference (Kabsch alignment), using the same $C_\alpha$ set for fitting and measurement (units: Å). Let $N_\alpha$ denote the number of $C_\alpha$ atoms with aligned positions $\{\mathbf{x}_i\}$ and reference positions $\{\mathbf{x}_i^{\text{ref}}\}$. Then

$$\text{RMSD} = \sqrt{\frac{1}{N_\alpha} \sum_{i=1}^{N_\alpha} \left\| \mathbf{x}_i - \mathbf{x}_i^{\text{ref}} \right\|^2}. \tag{S1-2}$$

The radius of gyration was computed over the same $C_\alpha$ set as

$$R_g = \sqrt{\frac{\sum_{i=1}^{N_\alpha} m_i \left\| \mathbf{x}_i - \mathbf{r}_{\text{cm}} \right\|^2}{\sum_{i=1}^{N_\alpha} m_i}}, \qquad \mathbf{r}_{\text{cm}} = \frac{\sum_{i=1}^{N_\alpha} m_i \mathbf{x}_i}{\sum_{i=1}^{N_\alpha} m_i}, \tag{S1-3}$$

which reduces to $R_g = \sqrt{\frac{1}{N_\alpha} \sum_i \|\mathbf{x}_i - \frac{1}{N_\alpha} \sum_j \mathbf{x}_j\|^2}$ when equal masses are assumed.

**Lag conversion.** (converted to frames) Given a desired lag $\tau$ (ns), the corresponding frame lag is

$$\ell \;=\; \mathrm{round}\Big(\frac{\tau}{\Delta t}\Big), \qquad \Delta t \;=\; \frac{\tau_T - \tau_1}{T-1}. \tag{S1-4}$$

#### S1.2 Minimum-barrier path construction and corridor selection

A transition corridor is defined in the selected collective-variable space. A discretized free-energy surface is first constructed, and a minimum-barrier path connecting the metastable basins is identified. The choice of this path is motivated by the relation $P(s) \propto \exp[-\beta F(s)]$, according to which lower-free-energy regions carry higher equilibrium probability. Thus, lower-barrier routes provide a natural approximation to statistically favorable transition channels in the projected landscape. A finite-width corridor is then constructed around the path, yielding a transition tube that isolates configurations near the putative transition region. This representation is computationally tractable, readily interpretable, and directly usable for subsequent mechanistic analysis. In future work, directional pathway selection could be refined by incorporating committor information or transition path theory (TPT)-based reactive fluxes, thereby prioritizing channels that are both statistically favorable in the free-energy landscape and dynamically important for reactive flow. Such an extension would require reliable Markovian models or transition-path ensembles from which committors and fluxes can be estimated, which is challenging for large systems with rare transitions.

In the present work, the two-dimensional free-energy surface is represented on a rectangular grid $\{(X_i, Y_j)\}$ with free-energy values $F_{ij}$. The user-specified start and end collective-variable coordinates, corresponding to the metastable states of interest, are mapped to the nearest grid nodes. A minimum-barrier path $\mathcal{P} = \{p_1, \ldots, p_L\}$ is then obtained on the grid, where $L = |\mathcal{P}|$ is the number of path nodes. Let $\mathbf{z}_\ell = (X_{p_\ell}, Y_{p_\ell})$, for $\ell = 1, \ldots, L$, denote the corresponding path coordinates. A finite corridor is defined around this path through geometric and free-energy tolerances, such that only configurations sufficiently close to the path and not excessively high in free energy are retained. Simulation frames are mapped to their nearest grid nodes in collective-variable space and retained only if those nodes lie within the corridor. The retained frames are then assigned to their nearest path positions, thereby defining the transition tube used in the subsequent analysis. In this construction, the path and corridor are defined directly from the free-energy landscape, yielding a direction-free progress coordinate along the transition tube.

A two-dimensional free-energy surface (FES) is represented on a rectangular grid $\{(X_i, Y_j)\}$ with free-energy values $F_{ij}$, expressed in units of $k_BT$. The user-specified start and end collective-variable coordinates are mapped to their nearest grid nodes, thereby defining the boundary nodes for the path construction. A minimum-barrier path

$$\mathcal{P} = \{p_1, \ldots, p_L\}$$

is then determined on the grid, where $L = |\mathcal{P}|$, with $p_1$ and $p_L$ denoting the snapped start and end nodes, respectively. Let the corresponding path-node coordinates be written as

$$\mathbf{z}_\ell = (X_{p_\ell}, Y_{p_\ell}), \qquad \ell = 1, \ldots, L,$$

with associated free-energy values

$$F(\mathbf{z}_\ell) = F_{p_\ell}.$$

A finite corridor around the path is subsequently introduced in order to select a transition-relevant subset of configuration space. At each path position $\ell$, the corridor is defined by a geometric radius $R_\ell$ in collective-variable units and a free-energy tolerance $T_\ell$ in units of $k_BT$. Simulation frames are mapped to their nearest grid nodes and are retained only if those nodes lie inside the corridor. The retained frames are then assigned to their nearest path positions and reported in chronological order, thereby defining the transition-path tube frames.

**Path on the free-energy grid.** On the FES grid, the optimal path is obtained by an A*-type search on the 8-connected grid, although 4-connectivity can also be used. The move cost is defined as a weighted combination of Euclidean step length and local free energy, together with a distance-based heuristic, so that both path length and barrier height are penalized. Specifically, the distance, free-energy, and heuristic terms are weighted by $(w_d, w_e, w_h)$, respectively. The search returns the ordered node sequence

$$\mathcal{P} = \{p_1, \ldots, p_L\}, \qquad p_1 = s, \quad p_L = e,$$

where $s$ and $e$ are the snapped start and end nodes. Optional postprocessing applies Gaussian smoothing to the path polyline in coordinate space with window size $w$ (disabled by default), after which the smoothed points are remapped to their nearest grid nodes.

**Corridor construction on the grid.** A corridor mask is constructed around the path by jointly constraining (i) the distance to the path in collective-variable space and (ii) the free-energy excess relative to the nearest path node. For each grid node $(i, j)$ with coordinates $(X_i, Y_j)$, the nearest path position is defined by

$$\ell^*(i,j) = \arg \min_{\ell \in \{1,\ldots,L\}} \left\|(X_i, Y_j) - \mathbf{z}_\ell\right\|_2. \tag{S1-5}$$

The corresponding distance to the path and free-energy excess are then given by

$$d_{ij} = \left\|(X_i, Y_j) - \mathbf{z}_{\ell^*(i,j)}\right\|_2, \tag{S1-6}$$

$$\Delta F_{ij} = F_{ij} - F_{p_{\ell^*(i,j)}}. \tag{S1-7}$$

Two path-indexed thresholds are defined: a local free-energy tolerance $T_\ell$ and a local maximum radius $R_\ell$. Both may be prescribed either as constants or through piecewise-linear interpolation along the path using the user parameters

$$(T^{(1)}_{\max}, T^{(2)}_{\max}, T_{\min}) \quad \text{and} \quad (R^{(1)}_{\max}, R^{(2)}_{\max}, R_{\min}),$$

together with a transition length $K \in \mathbb{N}$. Let

$$m = \ell - 1, \qquad n = L - \ell.$$

Then

$$T_\ell = \begin{cases} T^{(1)}_{\max} - \left(T^{(1)}_{\max} - T_{\min}\right)\dfrac{m}{K}, & m < K, \\ T^{(2)}_{\max} - \left(T^{(2)}_{\max} - T_{\min}\right)\dfrac{n}{K}, & n < K, \\ T_{\min}, & \text{otherwise}, \end{cases} \tag{S1-8}$$

and

$$R_\ell = \begin{cases} R^{(1)}_{\max} - \left(R^{(1)}_{\max} - R_{\min}\right)\dfrac{m}{K}, & m < K, \\ R^{(2)}_{\max} - \left(R^{(2)}_{\max} - R_{\min}\right)\dfrac{n}{K}, & n < K, \\ R_{\min}, & \text{otherwise}. \end{cases} \tag{S1-9}$$

Constant-width, constant-tolerance corridors are recovered when the end and interior values are chosen to be equal. A grid node $(i, j)$ is marked as inside the corridor if and only if

$$d_{ij} \leq R_{\ell^*(i,j)} \qquad \text{and} \qquad \Delta F_{ij} \leq T_{\ell^*(i,j)}. \tag{S1-10}$$

**Frame selection and ordering.** Each simulation frame with collective-variable coordinates $\mathbf{c}_t = (v^{(1)}_t, v^{(2)}_t)$ is mapped to its nearest grid node according to

$$\text{idx}(t) = \arg \min_{(i,j)} \left\|(X_i, Y_j) - \mathbf{c}_t\right\|_2. \tag{S1-11}$$

A frame is retained if and only if its nearest grid node lies inside the corridor mask. Retained frames are returned in ascending simulation index, i.e., in chronological order. By construction, each frame is included at most once.

**Default parameterization (as implemented).** For RMSD/$R_g$ and PCA-based CVs, constant thresholds are used:

$$T_{\max}^{(1)} = T_{\max}^{(2)} = T_{\min} = 0.20, \qquad R_{\max}^{(1)} = R_{\max}^{(2)} = R_{\min} = 0.20, \qquad K = 5.$$

For tICA-based CVs, end-specific thresholds tighter than the interior are used:

$$T_{\max}^{(1)} = T_{\max}^{(2)} = 0.15, \qquad T_{\min} = 0.20,$$

$$R_{\max}^{(1)} = R_{\max}^{(2)} = 0.15, \qquad R_{\min} = 0.20, \qquad K = 5.$$

In all cases, the path search uses 8-connectivity with

$$(w_d, w_e, w_h) = (1.0, 1.0, 1.0),$$

and smoothing is disabled by default.

**Summary.** A minimum-barrier path $\mathcal{P} = \{p_1, \ldots, p_L\}$ is obtained on the FES by an A*-type search with combined geometric and energetic costs. A corridor is then defined by path-indexed distance and free-energy tolerances $(R_\ell, T_\ell)$. Simulation frames are retained if their nearest grid nodes satisfy the corridor criteria, and the resulting transition-path tube frames are reported in chronological order.

*Feature set. PCA* Cartesian coordinates of atoms are used for principal component analysis (PCA) (after alignment) to ensure that retained components reflect large-amplitude, path-specific displacements. Exact atom selections and preprocessing (reference structure, alignment selection, units) are held fixed across all frames to maintain consistency.

#### S1.3 Causality: using corridor informativeness as a hypothesis generator

Corridor-conditioned informativeness is used here as a transition-focused screening criterion: it identifies players whose coordinates are strongly predictive of progress within the designated tube. Such predictiveness is valuable for mechanism discovery, but it is not, by itself, a causal statement: correlation structure and hidden confounding can yield strong association without direct causal influence.[1,2] Accordingly, causal relevance is not inferred from the ranking alone.

More broadly, purely state-wise uncertainty measures such as Shannon entropy $H(X)$ quantify uncertainty of a distribution, whereas transition control is inherently dynamical and can require time-directed information.[3] This distinction is explicit in Maxwell-demon and feedback-control settings, where thermodynamic performance depends on information acquired and used in a time-ordered control protocol.[4–6] Likewise, directed information-flow measures such as transfer entropy are designed to separate shared/static association from time-asymmetric influence by conditioning on history.[7,8]

A concrete example of directional leverage was observed in our earlier WalK histidine-kinase study, which aimed to induce a rare activation transition by steered MD.[9] In a reference protocol, an RMSD-based bias was applied to the C$\alpha$ atoms of both the DHp and CA domains to drive the transition. In an alternative protocol, the same bias was applied only to the DHp domain while the CA domain was left unperturbed. Strikingly, DHp-only driving produced an essentially comparable global rearrangement: DHp helical bending led to a concurrent CA rotation of $\sim 50°$, only $\sim 7°$ less than in the both-domain-biased protocol.[9] This illustrates that certain localized elements can carry transition-specific dynamical leverage over the collective rearrangement, motivating the present separation between corridor-informed screening (informativeness) and a subsequent causal analysis that targets directional influence on the designated transition. In the present framework, informativeness is therefore treated as a transition-focused hypothesis generator that nominates candidates, while causal support is sought separately by (i) robustness across distinct corridor constructions and (ii) by a dedicated intervention-tested causal analysis under controlled settings.

An equivalent, intuition-based view of the corridor-informativeness criterion is obtained by separating two requirements for a transition-relevant coordinate. First, it should be structurally expressed within the tube: if a coordinate remains nearly constant across corridor frames, then it cannot account for corridor progress except through rare, highly localized events. This requirement is quantified by the corridor-conditioned PCA variance, so that high PCA variance signals substantial transition-local rearrangement, whereas low variance signals an effectively stiff coordinate on the corridor. Second, the coordinate should be dynamically coherent with respect to slow progress: fluctuations that are large but fast, heterogeneous, or unstructured in time do not provide a reliable progress signal. This requirement is quantified by tICA participation, where high participation indicates alignment with the slow progress subspace along the time-ordered trajectory, whereas low participation indicates motion that does not track slow progress. The corridor-informativeness objective formalizes precisely this joint requirement by prioritizing coordinates that explain a substantial fraction of the variation of the retained slow progress coordinates on the corridor. The resulting PCA–tICA score therefore favors coordinates with both transition-local amplitude and slow dynamical coherence, while de-emphasizing stiff coordinates and fast idiosyncratic fluctuations.

At the ranking stage, redundancy is not explicitly penalized: players are first scored by their corridor-conditioned informativeness without imposing a global correlation-aware selection rule. This choice keeps the scoring step scalable, since evaluating all pairwise dependencies or optimizing a fully correlation-conditioned feature set would be computationally expensive for the high-dimensional structural representation used here. Redundancy is instead handled after ranking. Among the final selected candidates, strongly correlated players are identified using a correlation threshold, here $|\rho| > 0.7$. When two selected players exceed this threshold, only one representative is retained and the next highest-ranked non-redundant candidate is added. If correlated alternatives remain mechanistically plausible, the downstream controlled tests are repeated by substituting one candidate for the other, allowing their support to be assessed separately. This choice is deliberate: attempting to extract a minimal "causal feature set" from observational correlations requires additional causal assumptions and structure-learning machinery. [6]

For players, the resulting ranking is interpreted primarily as a measure of informativeness. Causal relevance is not inferred from the ranking alone, because dependence among candidate players and hidden confounding are not excluded by the score itself. Instead, causal support is sought separately from robustness across distinct CV constructions and from intervention-based verification.

Candidate selection then used consistent ranking across distinct global CV constructions, providing complementary views of the same transition. This reduced dependence on any single definition of the transition tube and helped expose transition-associated players that could be partially obscured in one projection. Causal support was then sought separately through intervention-based verification. When selected candidates were strongly correlated, here $|\rho| > 0.7$, only one representative was tested at a time, and correlated alternatives were substituted in separate controlled simulations to distinguish player-specific effects from redundant correlated reporters.

Independently of this post-ranking redundancy handling, causal support is sought downstream by triangulating evidence across two validation axes: (i) two separate controlled intervention-tested causal analyses and (ii) robustness to the three distinct global CV constructions used to define the transition tube.

#### S1.4 Consensus definition of the Hot-ROI.

For each protein, atoms were ranked independently within each of the three CV-defined corridors according to their corridor-conditioned informativeness score. The three corridors correspond to the RMSD/$R_g$, PCA-based, and tICA-based global CV constructions. Let $s_i^{(c)}$ denote the score of atom $i$ in corridor construction $c = 1, 2, 3$, and let $r_i^{(c)}$ denote its rank among the $M$ ranked atoms, with $r_i^{(c)} = 1$ corresponding to the largest score. For a target Hot-ROI size $N$, the top-$N$ atoms from each corridor were first selected, and atoms common to all three top-$N$ lists were retained.

If fewer than $N$ atoms were obtained, the rank depth was enlarged by two increments of approximately $N/3$, corresponding to an expanded depth of $N + 2\lceil N/3 \rceil$. Additional atoms appearing in the intersection of all three expanded lists were then added. This step was used only as a limited relaxation of the strict top-$N$ consensus, not as an unbounded expansion over all ranked atoms.

If the three-way consensus set still contained fewer than $N$ atoms, the remaining positions were filled using partial-consensus candidates, defined as atoms appearing in at least two of the three expanded high-rank lists. These candidates were ordered by a score-primary, rank-assisted criterion. To make scores comparable across corridor constructions, scores were normalized by the mean score within each corridor as $g_i^{(c)} = s_i^{(c)}/\mu_s^{(c)}$, where $\mu_s^{(c)}$ is the mean informativeness score over all atoms in corridor construction $c$. The aggregate normalized score was defined as $G_i = (1/3)\sum_{c=1}^{3} g_i^{(c)}$.

Ranks were also converted to percentile ranks as $p_i^{(c)} = 1 - (r_i^{(c)} - 1)/(M - 1)$, so that $p_i^{(c)} = 1$ for the highest-ranked atom and $p_i^{(c)} = 0$ for the lowest-ranked atom. The aggregate rank score was defined as $P_i = (1/3)\sum_{c=1}^{3} p_i^{(c)}$.

Partial-consensus candidates were ordered primarily by $G_i$. When two candidates differed by less than 5% in aggregate normalized score, namely when $|G_i - G_j|/\max(G_i, G_j) < 0.05$, the candidate with the larger $P_i$ was selected first. Candidates were added in this order until the Hot-ROI reached the target size $N$. Thus, the procedure first prioritizes atoms reproducibly high-ranked in all three CV-defined corridors, then fills any remaining positions from atoms supported by at least two corridors, using score magnitude as the primary criterion and rank consistency only to resolve near-ties.

#### S1.5 Interpretation and practical use

We propose a direct computational experiment for verification: by applying a reweighting enhanced sampling method like REST2 scaling-scheme *only* to the top-ranked player residues—effectively treating them as "hot spots"—one can test whether the free-energy barrier between the two metastable states is significantly reduced compared to a simulation where random or control residues are heated. A flattened, more accessible free-energy profile would provide compelling evidence that the identified players are not merely correlated with, but are functionally responsible for, governing the transition. This validation strategy, to our knowledge suggested here for the first time, moves beyond correlation to a more causal interpretation of mechanism. Progress will depend on analytical frameworks that convert computational and experimental structural ensembles into explicit, testable transition models, with predictive uncertainties quantified and concordance rigorously assessed against independent experimental and simulation observables.

### S2 Supporting Information 2

#### Mathematical framework for corridor-based informativeness and feature ranking

##### S2.1 State space, basins, and direction-free transition observable

Let $(X_t)_{t\geq 0}$ be a time-homogeneous stochastic process on a configuration/state space $\Omega$. Let $A, B \subset \Omega$ be two disjoint target sets, referred to as metastable basins.

The main formulation uses a direction-free transition observable

$$q : \Omega \to \mathbb{R}, \qquad x \mapsto q(x), \tag{S2-1}$$

to describe transition-relevant variation between the two basins.

A path or corridor of interest between the two basins is first selected, for example by high equilibrium probability, high reactive flux, a combined probability-flux criterion, or another physically motivated rule. Once this path ensemble has been specified, (q) is used passively to describe the structural and dynamical characteristics of the selected corridor. Thus, (q) does not define the path and is not required to have a global hitting-probability interpretation.

##### S2.2 corridor (tube/path) ensemble

All frames that lie in a corridor/tube region $\Theta \subset \Omega$ connecting $A$ and $B$ (e.g. a transition tube) are kept.

Denote the collected frames by

$$\mathcal{D}_\Theta := \{x^{(1)}, x^{(2)}, \ldots, x^{(N)}\} \subset \Theta.$$

The Method treats all frames equally (no reweighting) because long MD simulation is used.

$$\widehat{\mu}_\Theta := \frac{1}{N} \sum_{n=1}^{N} \delta_{x^{(n)}}. \tag{S2-2}$$

Equivalently, a random variable $X$ is defined by drawing a frame uniformly from $\mathcal{D}_\Theta$:

$$X \sim \widehat{\mu}_\Theta.$$

All expectations/variances/covariances/correlations in the method are taken with respect to $\widehat{\mu}_\Theta$ (over the corridor frames).

##### S2.3 Features and transition-progress coordinate

For each frame $x$, a feature vector is computed

$$\mathbf{X}(x) = (X_1(x), \ldots, X_d(x))^\top \in \mathbb{R}^d,$$

where each $X_i$ is a scalar feature (e.g. tied to an atom or an atomic descriptor). Let $S \subset \{1, \ldots, d\}$ be a subset of indices and $\mathbf{X}_S$ the subvector.

Consider a transition-progress coordinate $U$

$$U(x) := w^\top \mathbf{X}(x), \qquad w \in \mathbb{R}^d, \tag{S2-3}$$

where $w$ is a fixed loading vector (for example obtained from a kinetic model). The goal is to quantify how informative individual features, or subsets of features, are about the progress coordinate $U$.

#### S2.4 The selection objective (informativeness)

For a prescribed subset size $m$, the index set $S \subset 1, \dots, d$, with $|S| = m$, is selected to maximize the information retained by $\mathbf{X}_S$ about the transition observable $q(x)$ on the corridor-conditioned ensemble $\widehat{\mu}_\Theta$.

The informativeness of $S$ is defined as the explained-variance functional

$$I_\Theta(S) \ := \ \mathrm{Var}_{\widehat{\mu}_\Theta}\Big( \mathbb{E}_{\widehat{\mu}_\Theta}\big[q(X) \mid \mathbf{X}_S\big]\Big). \tag{S2-4}$$

Equivalently, by the law of total variance,

$$\mathrm{Var}_{\widehat{\mu}_\Theta}\big(q(X)\big) = \mathrm{Var}_{\widehat{\mu}_\Theta}\Big(\mathbb{E}_{\widehat{\mu}_\Theta}\big[q(X) \mid \mathbf{X}_S\big]\Big) + \mathbb{E}_{\widehat{\mu}_\Theta}\Big[\mathrm{Var}_{\widehat{\mu}_\Theta}\big(q(X) \mid \mathbf{X}_S\big)\Big]. \tag{S2-5}$$

Hence,

$$I_\Theta(S) = \mathrm{Var}_{\widehat{\mu}_\Theta}\big(q(X)\big) - \mathbb{E}_{\widehat{\mu}_\Theta}\Big[\mathrm{Var}_{\widehat{\mu}_\Theta}\big(q(X) \mid \mathbf{X}_S\big)\Big]. \tag{S2-6}$$

This shows that maximizing $I_\Theta(S)$ is equivalent to minimizing the unexplained conditional variance

$$\mathbb{E}_{\widehat{\mu}_\Theta}\Big[\mathrm{Var}_{\widehat{\mu}_\Theta}\big(q(X) \mid \mathbf{X}_S\big)\Big],$$

because the total variance $\mathrm{Var}_{\widehat{\mu}_\Theta}(q(X))$ does not depend on the choice of $S$.

Thus, $I_\Theta(S)$ is large when the selected features $\mathbf{X}_S$ account for a large fraction of the variation in the transition observable $q(x)$ across the corridor-conditioned ensemble, or equivalently when little uncertainty about the transition observable $q(x)$ remains after conditioning on $\mathbf{X}_S$. The value of $I_\Theta(S)$ therefore depends on both the chosen the transition observable $q(x)$ and the frame distribution $\widehat{\mu}_\Theta$.

#### S2.5 Covariance notation under the corridor ensemble

All moments in this subsection are taken with respect to the corridor-frame ensemble, that is, for $X \sim \widehat{\mu}_\Theta$.

The covariance matrix of the full feature vector is denoted by

$$\Sigma := \mathrm{Cov}_{\widehat{\mu}_\Theta}(\mathbf{X}) \in \mathbb{R}^{d\times d},$$

and the covariance matrix of the selected subvector $\mathbf{X}_S$ is denoted by

$$\Sigma_{SS} := \mathrm{Cov}_{\widehat{\mu}_\Theta}(\mathbf{X}_S) \in \mathbb{R}^{|S|\times|S|}.$$

The cross-covariance vector between the selected features and the scalar progress coordinate $U$ is defined by

$$\Sigma_{SU} := \mathrm{Cov}_{\widehat{\mu}_\Theta}(\mathbf{X}_S, U) \in \mathbb{R}^{|S|}.$$

Since

$$U = w^\top \mathbf{X},$$

the identity

$$\Sigma_{SU} = \mathrm{Cov}_{\widehat{\mu}_\Theta}(\mathbf{X}_S, w^\top \mathbf{X}) = \Sigma_{S\cdot}\, w \tag{S2-7}$$

is obtained by linearity of covariance, where $\Sigma_{S\cdot} \in \mathbb{R}^{|S|\times d}$ denotes the submatrix of $\Sigma$ formed by the rows indexed by $S$.

Thus, the vector $\Sigma_{SU}$ is fully determined by the pooled corridor covariance structure together with the loading vector $w$, and collects the covariances between the selected features and the progress coordinate.

##### S2.5.1 Covariance with a linear progress coordinate

Let the full feature vector be denoted by $X \in \mathbb{R}^d$, let $X_S$ denote the subvector indexed by $S$, and let the scalar progress coordinate be given by

$$U = w^\top X, \qquad w \in \mathbb{R}^d,$$

where $w$ is deterministic.

**Lemma 1** (Cross-covariance with a linear form)**.** *Let*

$$\Sigma := \mathrm{Cov}(X).$$

*Then*

$$\mathrm{Cov}(X_S, U) = \Sigma_{S\cdot} w,$$

*where $\Sigma_{S\cdot}$ denotes the submatrix of $\Sigma$ formed by the rows indexed by $S$.*

*Proof.* By definition,

$$\mathrm{Cov}(X_S, U) = \mathbb{E}[(X_S - \mathbb{E}[X_S])\,(U - \mathbb{E}[U])]\,.$$

Since $U = w^\top X$, it follows that

$$\mathbb{E}[U] = w^\top \mathbb{E}[X], \qquad U - \mathbb{E}[U] = w^\top (X - \mathbb{E}[X]).$$

Substitution therefore gives

$$\mathrm{Cov}(X_S, U) = \mathbb{E}\big[(X_S - \mathbb{E}[X_S])\, w^\top (X - \mathbb{E}[X])\big]\,.$$

By linearity, the vector $w$ may be taken outside the expectation, which yields

$$\mathrm{Cov}(X_S, U) = \mathbb{E}\big[(X_S - \mathbb{E}[X_S])(X - \mathbb{E}[X])^\top\big]\, w.$$

The expectation in brackets is precisely $\mathrm{Cov}(X_S, X)$, which is the row-block $\Sigma_{S\cdot}$ of $\Sigma$. Hence,

$$\mathrm{Cov}(X_S, U) = \Sigma_{S\cdot} w.$$

□

#### S2.6 Assumptions

The following assumptions are stated in a form under which the theorem below is exact. If one or more of them hold only approximately, then the resulting conclusion should also be interpreted as approximate.

(A1) **Pooled-frame corridor ensemble.** All expectations, variances, and covariances are taken with respect to $X \sim \widehat{\mu}_\Theta$, that is, the empirical distribution over the retained corridor frames. Since the analysis is based on long molecular-dynamics trajectories, this empirical corridor ensemble is taken to provide a representative sampling of the retained transition-tube configurations.

(A2) **Affine relation between transition observable and progress coordinate.** It is assumed that there exist constants $a \neq 0$ and $b \in \mathbb{R}$ such that

$$q(X) = a\,U + b$$

for $X \sim \widehat{\mu}_\Theta$.

(A3) **Affine conditional expectation of $U$ given $\mathbf{X}_S$.** It is assumed that the conditional expectation of $U$ given $\mathbf{X}_S$ is affine in $\mathbf{X}_S$, that is, that there exist $\alpha \in \mathbb{R}$ and $\beta_S \in \mathbb{R}^{|S|}$ such that

$$\mathbb{E}[U \mid \mathbf{X}_S] = \alpha + \beta_S^\top \mathbf{X}_S.$$

If the selected features are centered on the corridor ensemble, the intercept term is absorbed into the centering and the expression reduces to the linear form.

(A4) **Diagonal covariance within the selected coordinates.** It is assumed that, in the coordinate system used for ranking, the covariance of the selected features is diagonal, namely

$$\Sigma_{SS} = \mathrm{diag}(\sigma_i^2)_{i\in S}, \qquad \sigma_i^2 := \mathrm{Var}(X_i).$$

Under this assumption, the optimality criterion is reduced to a separable sum of per-feature contributions. The corresponding non-diagonal case, in which correlations between selected features are retained explicitly, is discussed in Sec. S2.13.

If this diagonal structure holds only approximately, then the resulting separable ranking should likewise be interpreted as an approximation.

In the present analysis, the diagonal form was used for the ranking step to avoid the computational cost of a fully correlation-conditioned selection. Redundancy was then controlled after ranking: when selected players showed strong correlation, here $|\rho| > 0.7$, only one representative was retained and the next-ranked candidate was included. This provides a computationally efficient approximation while avoiding simultaneous selection of strongly dependent players.

#### S2.7 Step 1: Reduction of transition-observable informativeness to progress-coordinate informativeness

Under assumption (A2), one has

$$\mathbb{E}\big[q(X) \mid \mathbf{X}_S\big] = \mathbb{E}\big[aU + b \mid \mathbf{X}_S\big] = a\,\mathbb{E}\big[U \mid \mathbf{X}_S\big] + b.$$

Taking the variance with respect to $\widehat{\mu}_\Theta$ eliminates the additive constant $b$ and yields

$$I_\Theta(S) = \mathrm{Var}_{\widehat{\mu}_\Theta}\Big(\mathbb{E}_{\widehat{\mu}_\Theta}\big[q(X) \mid \mathbf{X}_S\big]\Big) = a^2\,\mathrm{Var}_{\widehat{\mu}_\Theta}\Big(\mathbb{E}_{\widehat{\mu}_\Theta}\big[U \mid \mathbf{X}_S\big]\Big). \tag{S2-8}$$

Thus, under (A2), maximizing the informativeness of $\mathbf{X}_S$ about the transition observable $q$ is equivalent to maximizing the informativeness of $\mathbf{X}_S$ about the transition-progress coordinate $U$.

#### S2.8 Step 2: Closed-form expression for $\mathrm{Var}(\mathbb{E}[U \mid \mathbf{X}_S])$

Under assumption (A3), the conditional expectation of $U$ given $\mathbf{X}_S$ is affine,

$$\mathbb{E}[U \mid \mathbf{X}_S] = \alpha + \beta_S^\top \mathbf{X}_S, \tag{S2-9}$$

with $\alpha \in \mathbb{R}$ and $\beta_S \in \mathbb{R}^{|S|}$. Since $\alpha$ is constant, it does not contribute to the variance. Therefore,

$$\mathrm{Var}\Big(\mathbb{E}[U \mid \mathbf{X}_S]\Big) = \mathrm{Var}(\beta_S^\top \mathbf{X}_S) = \beta_S^\top \Sigma_{SS}\, \beta_S. \tag{S2-10}$$

For the affine least-squares regression of $U$ on $\mathbf{X}_S$, the coefficient vector is given by

$$\beta_S = \Sigma_{SS}^{-1}\Sigma_{SU}, \tag{S2-11}$$

provided that $\Sigma_{SS}$ is invertible. Substitution of Eq. (S2-11) into Eq. (S2-10) gives

$$\mathrm{Var}\Big(\mathbb{E}[U \mid \mathbf{X}_S]\Big) = \Sigma_{SU}^\top \Sigma_{SS}^{-1}\Sigma_{SU}. \tag{S2-12}$$

When this result is combined with Eq. (S2-9), the exact pooled-frame expression

$$I_\Theta(S) = a^2\, \Sigma_{SU}^\top \Sigma_{SS}^{-1}\Sigma_{SU} \tag{S2-13}$$

is obtained.

Using the identity $\Sigma_{SU} = \Sigma_{S\cdot} w$, this expression may also be written as

$$I_\Theta(S) = a^2\, w^\top \Sigma_{\cdot S}\, \Sigma_{SS}^{-1}\, \Sigma_{S\cdot}\, w, \tag{S2-14}$$

where $\Sigma_{\cdot S}$ denotes the submatrix of $\Sigma$ formed by the columns indexed by $S$.

**Critical note.** Equation (S2-14) is the correct general expression under assumptions (A1)–(A3). In the absence of (A4), the subset-selection problem is not, in general, reducible to a "top-$k$" rule based on independent per-feature scores, because the matrix $\Sigma_{SS}^{-1}$ couples the selected features and therefore accounts for redundancy among them.

##### S2.8.1 Variance of a affine form

Let $Z \in \mathbb{R}^m$ be a random vector with finite second moments, and let $\beta \in \mathbb{R}^m$ be deterministic. Then

$$\mathrm{Var}(\beta^\top Z) = \beta^\top \mathrm{Cov}(Z)\, \beta.$$

*Proof.* By definition,

$$\mathrm{Var}(\beta^\top Z) = \mathbb{E}\Big[\big(\beta^\top Z - \mathbb{E}[\beta^\top Z]\big)^2\Big].$$

By linearity of expectation,

$$\mathbb{E}[\beta^\top Z] = \beta^\top \mathbb{E}[Z],$$

and therefore

$$\beta^\top Z - \mathbb{E}[\beta^\top Z] = \beta^\top (Z - \mathbb{E}[Z]).$$

It follows that

$$\mathrm{Var}(\beta^\top Z) = \mathbb{E}\Big[\big(\beta^\top (Z - \mathbb{E}[Z])\big)^2\Big].$$

For any vector $a$, the identity

$$(\beta^\top a)^2 = \beta^\top (a a^\top) \beta$$

holds. Applying this with $a = Z - \mathbb{E}[Z]$ yields

$$\big(\beta^\top (Z - \mathbb{E}[Z])\big)^2 = \beta^\top \Big((Z - \mathbb{E}[Z])(Z - \mathbb{E}[Z])^\top\Big)\beta.$$

Taking expectations on both sides gives

$$\mathrm{Var}(\beta^\top Z) = \beta^\top\, \mathbb{E}\Big[(Z - \mathbb{E}[Z])(Z - \mathbb{E}[Z])^\top\Big]\, \beta.$$

The matrix inside the expectation is precisely the covariance matrix of $Z$, so

$$\mathrm{Var}(\beta^\top Z) = \beta^\top \mathrm{Cov}(Z)\, \beta.$$

□

##### S2.8.2 Derivation of $\beta_S = \Sigma_{SS}^{-1} \Sigma_{SU}$

Let $U$ be a scalar random variable with finite second moments, and let $X_S \in \mathbb{R}^m$ denote the selected feature subvector, where $m = |S|$. The residual is defined by

$$R := U - \mathbb{E}[U \mid X_S].$$

For any integrable function $g(X_S)$,

$$\mathbb{E}[R\, g(X_S)] = 0.$$

In particular,

$$\mathrm{Cov}(X_S, R) = 0.$$

*Proof.* By the tower property,

$$\mathbb{E}[R\, g(X_S)] = \mathbb{E}[\mathbb{E}[R\, g(X_S) \mid X_S]] = \mathbb{E}[g(X_S)\, \mathbb{E}[R \mid X_S]].$$

Since

$$\mathbb{E}[R \mid X_S] = \mathbb{E}[U - \mathbb{E}[U \mid X_S] \mid X_S] = 0,$$

the result follows. □

Now assume that the conditional expectation is affine in $X_S$, namely,

$$\mathbb{E}[U \mid X_S] = \alpha + \beta_S^\top X_S.$$

It then follows that

$$U = \alpha + \beta_S^\top X_S + R.$$

Taking the covariance with $X_S$ on both sides gives

$$\mathrm{Cov}(X_S, U) = \mathrm{Cov}(X_S, \alpha) + \mathrm{Cov}(X_S, \beta_S^\top X_S) + \mathrm{Cov}(X_S, R).$$

The first term vanishes because $\alpha$ is constant, and the third term vanishes by the orthogonality lemma. Hence,

$$\mathrm{Cov}(X_S, U) = \mathrm{Cov}(X_S, \beta_S^\top X_S).$$

Using the standard covariance identity for a linear form,

$$\mathrm{Cov}(X_S, \beta_S^\top X_S) = \mathrm{Cov}(X_S, X_S)\, \beta_S = \Sigma_{SS}\, \beta_S.$$

Therefore, the normal equations are obtained in the form

$$\Sigma_{SU} = \Sigma_{SS}\beta_S, \qquad \Sigma_{SU} := \mathrm{Cov}(X_S, U).$$

If $\Sigma_{SS}$ is invertible, this gives

$$\beta_S = \Sigma_{SS}^{-1}\Sigma_{SU}.$$

If $\Sigma_{SS}$ is singular, the corresponding least-squares solution is expressed in terms of the Moore–Penrose pseudoinverse as

$$\beta_S = \Sigma_{SS}^{\dagger}\Sigma_{SU}.$$

**Effect of subset-dependent omitted contributions.** The derivation above treats the selected representation $\mathbf{X}_S$ as the only set of variables entering the affine conditional model. In that case, the constant term $\alpha$ is fixed for the chosen subset $S$ and does not contribute to covariance. This is the assumption under which the normal equation $\Sigma_{SU} = \Sigma_{SS}\beta_S$ is obtained.

More generally, if the selected variables do not form a closed representation of the dependence of $U$, the contribution of the remaining variables may be written as a subset-dependent omitted component $A^{(S)}$. In this case,

$$U = \alpha + \beta_S^\top \mathbf{X}_S + A^{(S)} + R, \tag{S2-15}$$

where $A^{(S)}$ collects the part of the affine full-space predictor not represented explicitly by $\mathbf{X}_S$. Taking the covariance with $\mathbf{X}_S$ gives

$$\Sigma_{SU} = \Sigma_{SS}\beta_S + \Sigma_{SA}, \qquad \Sigma_{SA} := \mathrm{Cov}(\mathbf{X}_S, A^{(S)}), \tag{S2-16}$$

because the constant term does not contribute and the residual is orthogonal to the conditioning variables. Therefore, when $\Sigma_{SS}$ is invertible,

$$\beta_S = \Sigma_{SS}^{-1}\left(\Sigma_{SU} - \Sigma_{SA}\right). \tag{S2-17}$$

Thus, the simple expression $\beta_S = \Sigma_{SS}^{-1}\Sigma_{SU}$ is recovered only when the omitted component is uncorrelated with the selected variables, $\Sigma_{SA} = 0$, or when no such omitted component is present.

#### S2.9 Step 3: Separable form in the diagonal case

Under assumption (A4), the covariance matrix of the selected features is diagonal,

$$\Sigma_{SS} = \mathrm{diag}(\sigma_i^2)_{i\in S},$$

so that its inverse is also diagonal, with entries $1/\sigma_i^2$. Eq. (S2-13) then reduces to

$$I_\Theta(S) = a^2 \sum_{i\in S} \frac{\mathrm{Cov}(X_i, U)^2}{\mathrm{Var}(X_i)}. \tag{S2-18}$$

A separable sum of nonnegative per-feature contributions is therefore obtained.

#### S2.10 Step 4: Rewriting as variance times squared correlation

The correlation between $X_i$ and $U$ is defined by

$$\mathrm{Corr}(X_i, U) := \frac{\mathrm{Cov}(X_i, U)}{\sqrt{\mathrm{Var}(X_i)\,\mathrm{Var}(U)}}.$$

It follows immediately that

$$\frac{\mathrm{Cov}(X_i, U)^2}{\mathrm{Var}(X_i)} = \mathrm{Var}(U)\,\mathrm{Corr}(X_i, U)^2. \tag{S2-19}$$

Substitution of Eq. (S2-19) into Eq. (S2-18) yields

$$I_\Theta(S) = a^2\,\mathrm{Var}(U) \sum_{i \in S} \mathrm{Corr}(X_i, U)^2. \tag{S2-20}$$

Since the prefactor $a^2\,\mathrm{Var}(U)$ does not depend on the choice of $S$, maximization of $I_\Theta(S)$ over all subsets of fixed size $|S| = k$ is equivalent to maximization of

$$\sum_{i \in S} s_i, \qquad s_i := \mathrm{Corr}(X_i, U)^2. \tag{S2-21}$$

Thus, in the diagonal case, the selection criterion is reduced to a sum of independent per-feature scores, and the optimal subset of size $k$ is obtained by choosing the $k$ features with the largest values of $s_i$.

#### S2.11 Theorem ( statement for the pooled-frame method)

**Theorem 1** (Optimality of the "variance $\times$ slow-loading-squared" rule under pooled frames)**.** *Assume (A1)–(A4), and fix an integer $k \in \{1, \ldots, d\}$. Let the per-feature scores $s_i$ be defined by Eq.* (S2-21)*. Then any maximizer*

$$S^\star \in \arg \max_{S \subset \{1,\ldots,d\},\ |S|=k} I_\Theta(S)$$

*may be chosen as the set of the $k$ indices with the largest values of $s_i$. Equivalently, selection of the top-$k$ features according to $s_i$ maximizes $I_\Theta(S)$.*

*Proof.* Under assumptions (A1)–(A4), Eq. (S2-20) gives

$$I_\Theta(S) = C \sum_{i \in S} s_i, \qquad C := a^2\,\mathrm{Var}(U) > 0.$$

For fixed $k$, maximization of $I_\Theta(S)$ is therefore equivalent to maximization of

$$\sum_{i \in S} s_i$$

over all subsets $S$ with $|S| = k$. Since this objective is separable and each selected feature contributes one unit to the cardinality constraint, an optimal subset is obtained by selecting the $k$ largest scores $s_i$. □

#### S2.12 Connection to the "slow loading" form in the globally diagonal case

If, in addition, the full covariance matrix $\Sigma$ is diagonal, so that the working coordinates are mutually uncorrelated, then for each $i$ one has

$$\mathrm{Cov}(X_i, U) = \mathrm{Cov}\Big(X_i, \sum_{j=1}^{d} w_j X_j\Big) = \sum_{j=1}^{d} w_j\,\mathrm{Cov}(X_i, X_j) = w_i\,\mathrm{Var}(X_i). \tag{S2-22}$$

Substitution of Eq. (S2-22) into Eq. (S2-18) yields

$$\frac{\mathrm{Cov}(X_i, U)^2}{\mathrm{Var}(X_i)} = \frac{\big(w_i\,\mathrm{Var}(X_i)\big)^2}{\mathrm{Var}(X_i)} = w_i^2\,\mathrm{Var}(X_i). \tag{S2-23}$$

Accordingly, in the globally diagonal case,

$$I_\Theta(S) = a^2 \sum_{i \in S} w_i^2 \operatorname{Var}(X_i), \tag{S2-24}$$

and the per-feature score reduces to

$$s_i = w_i^2 \operatorname{Var}(X_i). \tag{S2-25}$$

Thus, in this special case, the optimal ranking is exactly the "variance $\times$ slow-loading-squared" rule.

**(1) Exactness hinges on assumptions (A2)–(A4).** If the transition observable is not exactly affine in $U$ on $\Theta$, if $\mathbb{E}[U \mid \mathbf{X}_S]$ is not affine in $\mathbf{X}_S$, or if $\Sigma_{SS}$ is not diagonal, then the top-$k$ rule is no longer guaranteed to be optimal for $I_\Theta(S)$. This is the best linear approximation. In particular, when $\Sigma_{SS}$ contains substantial off-diagonal correlations, the selected features are coupled through $\Sigma_{SS}^{-1}$, and the true optimum depends on the subset $S$ itself through Eqs. (S2-13)–(S2-14) which is discussed in Sec. S2.13.

#### S2.13 Non-diagonal $\Sigma_{SS}$: exact set-level informativeness in correlation–variance form

##### S2.13.1 Exact set-level informativeness under affine least-squares regression

The affine transition observable $q(x)$ relation on the corridor is retained in the form

$$q(X) = aU + b, \qquad a \neq 0. \tag{S2-26}$$

An affine model is also assumed for the conditional expectation of $U$ given $\mathbf{X}_S$.

Under these assumptions,

$$I_\Theta(S) = \operatorname{Var}\big(\mathbb{E}[q(X) \mid X_S]\big) = a^2 \operatorname{Var}\big(\mathbb{E}[U \mid X_S]\big). \tag{S2-27}$$

Without any diagonality assumption, the exact expression

$$\operatorname{Var}\big(\mathbb{E}[U \mid X_S]\big) = \Sigma_{SU}^\top \Sigma_{SS}^{-1} \Sigma_{SU} \tag{S2-28}$$

is obtained, and therefore

$$I_\Theta(S) = a^2\, \Sigma_{SU}^\top \Sigma_{SS}^{-1} \Sigma_{SU}. \tag{S2-29}$$

##### S2.13.2 Rewriting in terms of variances and correlations

Let $m = |S|$, and define

$$\sigma_i := \sqrt{\operatorname{Var}(X_i)} \qquad (i \in S), \qquad \sigma_U := \sqrt{\operatorname{Var}(U)}. \tag{S2-30}$$

Let

$$D := \operatorname{diag}(\sigma_1, \ldots, \sigma_m). \tag{S2-31}$$

Furthermore, let

$$R := \operatorname{Corr}(X_S) \in \mathbb{R}^{m \times m}, \qquad r := \operatorname{Corr}(X_S, U) \in \mathbb{R}^m, \tag{S2-32}$$

so that

$$R_{ij} = \operatorname{Corr}(X_i, X_j), \qquad r_i = \operatorname{Corr}(X_i, U). \tag{S2-33}$$

With this notation, the covariance factorizations

$$\Sigma_{SS} = DRD, \qquad \Sigma_{SU} = \sigma_U\, Dr \tag{S2-34}$$

are exact. If $R$ is invertible, equivalently if $\Sigma_{SS}$ is invertible, then

$$\Sigma_{SS}^{-1} = D^{-1} R^{-1} D^{-1}. \tag{S2-35}$$

Substitution of Eqs. (S2-34) and (S2-35) into Eq. (S2-28) gives

$$\operatorname{Var}\big(\mathbb{E}[U \mid X_S]\big) = \sigma_U^2\, r^\top R^{-1} r = \operatorname{Var}(U)\, r^\top R^{-1} r. \tag{S2-36}$$

Hence,

$$I_\Theta(S) = a^2 \operatorname{Var}(U)\, r^\top R^{-1} r. \tag{S2-37}$$

If $R$ is singular, $R^{-1}$ is replaced by the Moore–Penrose pseudoinverse $R^\dagger$.

**Interpretation.** The scalar $r^\top R^{-1} r$ is the squared multiple correlation, that is, the linear-regression $R^2$ of $U$ on $X_S$ after redundancy among the selected features has been taken into account.

###### S2.13.3 Diagonal special case as a corollary

If the selected features are uncorrelated under $\widehat{\mu}_\Theta$, then $R = I$, and therefore

$$r^\top R^{-1} r = r^\top r = \sum_{i \in S} \operatorname{Corr}(X_i, U)^2. \tag{S2-38}$$

It follows that

$$\operatorname{Var}\big(\mathbb{E}[U \mid X_S]\big) = \operatorname{Var}(U) \sum_{i \in S} \operatorname{Corr}(X_i, U)^2. \tag{S2-39}$$

This is the separable case in which the objective decomposes into additive per-feature contributions, so that top-$k$ selection by an individual feature score becomes valid.

###### S2.13.4 Where the approximation enters

The identity relating covariance, variance, and correlation is exact:

$$\frac{\operatorname{Cov}(Z, U)^2}{\operatorname{Var}(Z)} = \operatorname{Var}(U) \operatorname{Corr}(Z, U)^2. \tag{S2-40}$$

The approximation in the "variance × loading" rule enters only when the exact quantity

$$r^\top R^{-1} r \tag{S2-41}$$

is replaced by

$$r^\top r, \tag{S2-42}$$

that is, when

$$R \approx I \qquad \Longrightarrow \qquad R^{-1} \approx I \tag{S2-43}$$

is assumed.

A first-order expansion may be written by setting

$$R = I + E, \tag{S2-44}$$

where $E$ contains the off-diagonal correlations and is assumed to be small. Then

$$R^{-1} = (I + E)^{-1} \approx I - E, \tag{S2-45}$$

which yields

$$r^\top R^{-1} r \approx r^\top r - r^\top E r. \tag{S2-46}$$

Thus, positive correlations among features generally reduce the marginal gain obtained by selecting multiple similar features, reflecting redundancy within the selected set.

##### S2.14 Two-dimensional slow coordinate: $U \in \mathbb{R}^2$

###### S2.14.1 Definition of the two-dimensional target and its conditional mean

Let the slow target be the two-dimensional random vector

$$U(X) = \begin{pmatrix} U^{(1)}(X) \\ U^{(2)}(X) \end{pmatrix} \in \mathbb{R}^2, \tag{S2-47}$$

for example $(\mathrm{tIC}_1, \mathrm{tIC}_2)$ obtained from a tICA computation at lag $\tau$. Let $X_S \in \mathbb{R}^m$ denote the selected feature subvector, where $m = |S|$. The associated conditional mean predictor is

$$\mathbb{E}[U \mid X_S] \in \mathbb{R}^2. \tag{S2-48}$$

##### S2.14.2 Matrix-valued informativeness: explained covariance

The direct multivariate analogue of $\mathrm{Var}(\mathbb{E}[U \mid X_S])$ is the explained covariance matrix

$$\mathcal{I}(S) := \mathrm{Cov}\big(\mathbb{E}[U \mid X_S]\big) \in \mathbb{R}^{2\times 2}. \tag{S2-49}$$

This matrix quantifies how much of the variance of each component of $U$, together with their cross-covariance, is explained by the selected features $X_S$.

##### S2.14.3 Scalar summaries of informativeness

To compare different subsets $S$ by a single scalar quantity, the trace may be used:

$$I_{\mathrm{tr}}(S) := \operatorname{tr} \mathcal{I}(S). \tag{S2-50}$$

This equals the total explained variance in $U^{(1)}$ and $U^{(2)}$.

##### S2.14.4 Closed form in terms of covariance blocks

All covariance matrices in this subsection are taken under the same frame ensemble, as in the preceding one-dimensional derivation. Define

$$\Sigma_{SS} := \mathrm{Cov}(X_S) \in \mathbb{R}^{m\times m}, \qquad \Sigma_{SU} := \mathrm{Cov}(X_S, U) \in \mathbb{R}^{m\times 2}, \tag{S2-51}$$

and let

$$\Sigma_{US} := \Sigma_{SU}^{\top} \in \mathbb{R}^{2\times m}. \tag{S2-52}$$

Under the same affine-projection interpretation used in the one-dimensional case (exact when $(\mathbf{X}_S, U)$ is jointly Gaussian, and otherwise interpreted in terms of the best affine predictor), the explained covariance is given by

$$\mathcal{I}(S) = \mathrm{Cov}\big(\mathbb{E}_{\mathrm{aff}}[U \mid \mathbf{X}_S]\big) = \Sigma_{US}\, \Sigma_{SS}^{-1}\, \Sigma_{SU}. \tag{S2-53}$$

Accordingly, the trace-based informativeness is

$$I_{\mathrm{tr}}(S) = \operatorname{tr}\big(\Sigma_{US}\, \Sigma_{SS}^{-1}\, \Sigma_{SU}\big)\,, \tag{S2-54}$$

If $\Sigma_{SS}$ is singular, $\Sigma_{SS}^{-1}$ is replaced by the Moore–Penrose pseudoinverse $\Sigma_{SS}^{\dagger}$.

##### S2.14.5 Diagonal special case and additive per-feature scores

If the selected features are uncorrelated in the working basis, so that $\Sigma_{SS}$ is diagonal with entries $\mathrm{Var}(X_i)$, then $\Sigma_{SS}^{-1}$ is also diagonal and the trace criterion becomes additive:

$$I_{\mathrm{tr}}(S) = \sum_{i\in S} \sum_{k=1}^{2} \frac{\mathrm{Cov}(X_i, U^{(k)})^2}{\mathrm{Var}(X_i)}. \tag{S2-55}$$

A natural per-feature score is therefore

$$s_i := \sum_{k=1}^{2} \frac{\mathrm{Cov}(X_i, U^{(k)})^2}{\mathrm{Var}(X_i)}. \tag{S2-56}$$

Using the identity

$$\frac{\mathrm{Cov}(X_i, U^{(k)})^2}{\mathrm{Var}(X_i)} = \mathrm{Var}(U^{(k)})\, \mathrm{Corr}(X_i, U^{(k)})^2, \tag{S2-57}$$

the same score may be written as

$$s_i = \sum_{k=1}^{2} \mathrm{Var}(U^{(k)})\, \mathrm{Corr}(X_i, U^{(k)})^2. \tag{S2-58}$$

Since $\mathrm{Var}(U^{(1)})$ and $\mathrm{Var}(U^{(2)})$ do not depend on the feature index $i$, they act only as global component weights in the ranking.

##### S2.14.6 Specialization to PCA and tICA coordinates

Let $Y \in \mathbb{R}^m$ denote PCA scores in a basis for which

$$\mathrm{Cov}(Y) = \Lambda = \mathrm{diag}(\lambda_1, \ldots, \lambda_m). \tag{S2-59}$$

Suppose that the two slow coordinates are represented in PCA-score coordinates as

$$U^{(1)} = a^{(1)\top} Y, \qquad U^{(2)} = a^{(2)\top} Y, \tag{S2-60}$$

with coefficient vectors $a^{(1)}, a^{(2)} \in \mathbb{R}^m$.

Then, by the diagonal-case formula above,

$$\mathrm{Cov}(Y_i, U^{(1)}) = \lambda_i a_i^{(1)}, \qquad \mathrm{Cov}(Y_i, U^{(2)}) = \lambda_i a_i^{(2)}, \tag{S2-61}$$

and the induced per-coordinate score becomes

$$s_i = \lambda_i \Big( (a_i^{(1)})^2 + (a_i^{(2)})^2 \Big). \tag{S2-62}$$

#### S2.15 General-dimensional slow coordinate: $U \in \mathbb{R}^r$

##### S2.15.1 Definition of the $r$-dimensional target and its conditional mean

Let the slow target be the $r$-dimensional random vector

$$U(X) = \begin{pmatrix} U^{(1)}(X) \\ \vdots \\ U^{(r)}(X) \end{pmatrix} \in \mathbb{R}^r, \tag{S2-63}$$

for example $(\mathrm{tIC}_1, \ldots, \mathrm{tIC}_r)$ obtained from a tICA computation at lag $\tau$. Let $X_S \in \mathbb{R}^m$ denote the selected feature subvector, where $m = |S|$. The corresponding conditional mean predictor is

$$\mathbb{E}[U \mid X_S] \in \mathbb{R}^r. \tag{S2-64}$$

##### S2.15.2 Matrix-valued informativeness: explained covariance

The direct multivariate analogue of the one-dimensional explained variance is the explained covariance matrix

$$\mathcal{I}(S) := \mathrm{Cov}\big(\mathbb{E}[U \mid X_S]\big) \in \mathbb{R}^{r \times r}. \tag{S2-65}$$

This matrix quantifies how much of the variance of each component of $U$, together with the cross-covariances among components, is explained by the selected features $X_S$.

##### S2.15.3 Scalar summaries of informativeness

To compare different subsets $S$ by a single scalar quantity, the trace may be used:

$$I_{\mathrm{tr}}(S) := \mathrm{tr}\, \mathcal{I}(S). \tag{S2-66}$$

This equals the total explained variance across the $r$ components of $U$.

##### S2.15.4 Closed form in terms of covariance blocks

All covariance matrices in this subsection are taken under the same frame ensemble as in the preceding derivations. Define

$$\Sigma_{SS} := \mathrm{Cov}(X_S) \in \mathbb{R}^{m \times m}, \qquad \Sigma_{SU} := \mathrm{Cov}(X_S, U) \in \mathbb{R}^{m \times r}, \tag{S2-67}$$

and let

$$\Sigma_{US} := \Sigma_{SU}^{\top} \in \mathbb{R}^{r\times m}. \tag{S2-68}$$

If the conditional mean is exactly affine in $X_S$, then the explained covariance is given by

$$\mathcal{I}(S) = \mathrm{Cov}\big(\mathbb{E}[U \mid X_S]\big) = \Sigma_{US}\,\Sigma_{SS}^{-1}\,\Sigma_{SU}. \tag{S2-69}$$

This holds, for example, when $(X_S, U)$ is jointly Gaussian.

More generally, even when the conditional mean is not exactly affine, the matrix

$$\Sigma_{US}\,\Sigma_{SS}^{-1}\,\Sigma_{SU} \tag{S2-70}$$

is still the covariance explained by the best affine predictor of $U$ from $\mathbf{X}_S$. Accordingly, the trace-based informativeness criteria is

$$I_{\mathrm{tr}}^{\mathrm{lin}}(S) = \mathrm{tr}\left(\Sigma_{US}\,\Sigma_{SS}^{-1}\,\Sigma_{SU}\right), \tag{S2-71}$$

If $\Sigma_{SS}$ is singular, $\Sigma_{SS}^{-1}$ is replaced by the Moore–Penrose pseudoinverse $\Sigma_{SS}^{\dagger}$.

##### S2.15.5 Diagonal special case and additive per-feature scores

If the selected features are uncorrelated in the working basis, so that $\Sigma_{SS}$ is diagonal with entries $\mathrm{Var}(X_i)$, then the trace criterion becomes additive across features:

$$I_{\mathrm{tr}}^{\mathrm{lin}}(S) = \sum_{i\in S} \underbrace{\sum_{k=1}^{r} \frac{\mathrm{Cov}(X_i, U^{(k)})^2}{\mathrm{Var}(X_i)}}_{=:s_i}. \tag{S2-72}$$

A natural per-feature score is therefore

$$s_i = \sum_{k=1}^{r} \frac{\mathrm{Cov}(X_i, U^{(k)})^2}{\mathrm{Var}(X_i)}. \tag{S2-73}$$

Using the identity

$$\frac{\mathrm{Cov}(X_i, U^{(k)})^2}{\mathrm{Var}(X_i)} = \mathrm{Var}(U^{(k)})\,\mathrm{Corr}(X_i, U^{(k)})^2, \tag{S2-74}$$

the same score may be written as

$$s_i = \sum_{k=1}^{r} \mathrm{Var}(U^{(k)})\,\mathrm{Corr}(X_i, U^{(k)})^2. \tag{S2-75}$$

This expression is exact.

##### S2.15.6 Specialization to PCA and tICA coordinates

Let $Y \in \mathbb{R}^m$ denote PCA scores in a basis for which

$$\mathrm{Cov}(Y) = \Lambda = \mathrm{diag}(\lambda_1, \ldots, \lambda_m). \tag{S2-76}$$

Assume that the $r$ slow coordinates are represented in PCA-score coordinates as

$$U^{(k)} = a^{(k)\top} Y, \qquad k = 1, \ldots, r, \tag{S2-77}$$

with coefficient vectors $a^{(k)} \in \mathbb{R}^m$.

Then

$$\mathrm{Cov}(Y_i, U^{(k)}) = \lambda_i a_i^{(k)}, \tag{S2-78}$$

and the induced per-coordinate score becomes

$$s_i = \lambda_i \sum_{k=1}^{r} (a_i^{(k)})^2. \tag{S2-79}$$

##### S2.15.7 Interpretation

For $r = 1$, this construction reduces to the one-dimensional explained-variance criterion. For $r > 1$, $\mathcal{I}(S)$ measures how well $X_S$ explains the slow subspace spanned by $(U^{(1)}, \dots, U^{(r)})$.

When $\operatorname{tr}\mathcal{I}(S)$, or its linear-predictor analogue $I^{\mathrm{lin}}_{\mathrm{tr}}(S)$, is used, subsets are favored that explain variance in any component of the slow target. The resulting selected features may therefore be more diverse than in the one-dimensional case, because multiple slow directions are rewarded rather than a single direction alone.

#### S2.16 Consequence for the ranking formula, exact non-diagonal form, and practical scope

The preceding derivations show that the ranking rule used in the present work can be placed on a precise mathematical footing.

For a one-dimensional slow target, the exact additive score in a diagonal ranking basis is

$$s_i = \operatorname{Var}(Y_i)\left(a_i^{(1)}\right)^2. \tag{S2-80}$$

For an $r$-dimensional slow target

$$U = (U^{(1)}, \dots, U^{(r)}), \qquad U^{(k)} = a^{(k)\top} Y, \tag{S2-81}$$

the exact trace-based multivariate generalization is

$$s_i = \operatorname{Var}(Y_i) \sum_{k=1}^{r} \left(a_i^{(k)}\right)^2. \tag{S2-82}$$

In the PCA score basis, where

$$\operatorname{Cov}(Y) = \Lambda = \operatorname{diag}(\lambda_1, \dots, \lambda_m), \tag{S2-83}$$

this becomes

$$s_i = \lambda_i \sum_{k=1}^{r} \left(a_i^{(k)}\right)^2. \tag{S2-84}$$

Thus, the rule "variance times sum of squared tICA loadings" is not merely heuristic: it is obtained exactly from the multivariate explained-covariance objective when the assumptions of the derivation hold.

More specifically, under the linear-predictor informativeness criterion

$$I^{\mathrm{lin}}_{\mathrm{tr}}(S) = \operatorname{tr}\left(\Sigma_{US}\Sigma_{SS}^{-1}\Sigma_{SU}\right), \tag{S2-85}$$

and under the diagonal-covariance assumption in the ranking basis, the subset-selection problem with fixed cardinality $|S| = k$ reduces to

$$I^{\mathrm{lin}}_{\mathrm{tr}}(S) = \sum_{i \in S} s_i, \tag{S2-86}$$

with $s_i$ given by Eq. (S2-82). It then follows that an optimal subset is obtained by selecting the $k$ features with the largest values of $s_i$.

**Exact treatment when feature correlations are non-negligible.** If the selected features are correlated, then the diagonal reduction is no longer exact. In that case, the exact set-level objective has already been obtained in the form

$$I^{\mathrm{lin}}_{\mathrm{tr}}(S) = \operatorname{tr}\left(\Sigma_{US}\Sigma_{SS}^{-1}\Sigma_{SU}\right), \tag{S2-87}$$

or, equivalently, in correlation–redundancy form,

$$I^{\mathrm{lin}}_{\mathrm{tr}}(S) = \operatorname{tr}\left(W_U\, R_{UX_S} R_{X_S X_S}^{-1} R_{X_S U}\right) \tag{S2-88}$$

for a suitable normalization of the target, with the simpler scalar expression $r^\top R^{-1} r$ arising in the one-dimensional case. The matrix $\Sigma_{SS}^{-1}$ explicitly accounts for redundancy among selected features. Therefore, when off-diagonal correlations are appreciable, the exact optimum is set-dependent, and the top-$k$ rule based on independent per-feature scores should be understood only as an approximation.

**Why the assumptions are reasonable in practice.** The present derivation is formulated at the level of the best linear approximation. This choice is intentional: it makes the mechanism of the ranking transparent and yields an objective that can be analyzed exactly. More flexible nonlinear extensions, including machine-learning models, could be developed in future work. Nevertheless, the assumptions used here are not ad hoc; in the present setting they are reasonable for the following practical and empirical reasons.

The exact derivation requires a sequence of structural assumptions. These assumptions are restrictive, but they are not artificial, and in the present setting they can be made plausible.

First, the use of the corridor-frame ensemble is natural because the scientific objective is not to explain equilibrium variation globally, but to explain variation relevant to the designated transition event. If the corridor is constructed from a collective-variable representation that resolves the dominant transition channel well, then the retained frames form a transition-enriched ensemble on which a progress-oriented ranking criterion is meaningful.

Second, the reduction from the transition-observable informativeness to slow-coordinate informativeness is reasonable when the chosen slow coordinate, or slow subspace, is a good surrogate for progress along the transition. This does not require the transition observable to be known exactly. It only requires that, on the retained corridor frames, the transition observable be well approximated by an affine function of the selected slow coordinate(s). If the corridor has been constructed from informative collective variables and the transition is effectively low-dimensional in that representation, this approximation can be sufficiently accurate to justify the reduction operationally.

Third, the linear-predictor step is exact when the joint distribution is Gaussian, but it remains meaningful more generally because it defines the best linear predictor of the slow target from the selected features. In this sense, the derivation identifies the exact optimizer of a linear informativeness objective even when the full conditional mean is nonlinear. This is often a useful approximation in practice, especially after dimensionality reduction and restriction to a narrow corridor.

Fourth, the diagonal-covariance assumption should not be imposed blindly, but should be checked for the coordinates used in the ranking. If the selected features are only weakly dependent under the working ensemble — for example, if the pairwise absolute correlations are generally small, according to a practical threshold chosen by the analyst — then the diagonal approximation may be used and the separable score provides a reasonable description. By contrast, if substantial dependence remains among the selected features, then the exact non-diagonal formula should be used, since redundancy enters through $\Sigma_{SS}^{-1}$. In practice, this also motivates the use of alternative collective-variable representations, such as PCA coordinates or other more decorrelated bases, when the original features do not satisfy the approximation sufficiently well.

**When the assumptions are only partially satisfied.** If the selected features, the chosen slow coordinates, or the working basis do not satisfy the assumptions sufficiently well, then two routes remain available.

The first is to retain the more exact set-level formula, Eq. (S2-87), and to perform a more specific analysis that accounts explicitly for correlation and redundancy among features. This is the mathematically preferred route when strong off-diagonal structure is present.

The second is to accept the diagonal reduction as a controlled approximation. In that case, the score in Eq. (S2-84) should be interpreted as a principled surrogate that is expected to work well only insofar as the chosen basis approximately decorrelates the relevant features and the chosen slow coordinates capture the transition progress with sufficient fidelity.

**Possible extensions beyond the present linear framework.** The present derivation is intentionally linear. It therefore clarifies exactly where the assumptions enter, but it also indicates natural directions for further improvement.

In particular, machine-learning approaches could be used in future work to address two of the present limitations. First, they could assist in the construction of more informative collective variables or slow coordinates, thereby improving the compatibility of the corridor and target with the assumptions underlying the derivation. Second, nonlinear predictors could be introduced in place of the linear conditional-mean model, allowing a more flexible notion of informativeness than the present linear-predictor criterion. Such extensions would not invalidate the current derivation; rather, they would generalize it beyond the linear regime considered here.

**Applicability to both motions and players.** The informativeness framework introduced above is sufficiently general to apply to both levels of analysis considered in the present work.

At the level of *motions*, the collective variables are constructed by the corridor-conditioned PCA–tICA procedure itself. PCA is first performed on the corridor frames in order to define a transition-focused representation. The trajectory is then projected into this corridor-defined subspace, and slow collective variables are obtained by a weighted tICA step written in generalized form, so that corridor-relevant frames or lag pairs can be emphasized directly in the dynamical estimator. In this way, the final collective variables are not generic global slow modes, but corridor-focused slow coordinates designed to capture the dominant transition-relevant directions within the selected representation. The resulting motion ranking should therefore be interpreted as quantifying how strongly each motion contributes to these weighted, corridor conditioned slow collective variables.

At the level of *players*, the variables $X_i$ may instead be interpreted as player-specific descriptors, grouped coordinates, or atom-/residue-level contributions. In that setting, the same objective quantifies how informative each player, or each subset of players, is about the slow transition coordinates. The mathematical criterion is unchanged; only the interpretation of the features differs.

**Practical conclusion.** The derivation therefore supports the following statement.

If a good corridor has been constructed, if the retained slow coordinates provide a faithful local representation of transition progress, and if the ranking is performed in a decorrelated basis, then the score

$$s_i = \lambda_i \sum_{k=1}^{r} \left(a_i^{(k)}\right)^2 \tag{S2-89}$$

is not only reasonable but is obtained exactly as the optimizer of the corresponding multivariate explained-covariance criterion.

When these assumptions are not fully satisfied, the exact correlation-aware formula remains available, whereas the simpler product score should be interpreted as a principled approximation. In this sense, the present framework provides both an exact idealized justification and a practical working rule for ranking informative motions or players.

Table S3-1: ROI cohorts used for validation

| Cohort class | ROI definition |
|---|---|
| Hot-ROI | Top-$N$ atoms ranked by the corridor-conditioned score $S_\alpha$. |
| Random-ROI | Randomly selected $N$ atoms. |
| Tail-ROI | Top $M$ atoms by $S_\alpha$ were removed; the next $N$ ranked atoms were used. |
| Ranking ablations | Path-PCA-ROI, Path-tICA-ROI, and G-tICA-ROI: atoms ranked using PCA-only, path-tICA-only, or global tICA. |
| Data-fraction test | Scores recomputed using different percentages of the reference trajectory. |
| Metadynamics test | Same CV and multi-walker well-tempered settings; biased atom set varied among all-atoms, Hot-ROI, and Random-ROI. |

$N$ and $M$ were matched across cohorts per protein. REST2 scaling schedules and simulation time budgets were held identical across matched ensembles.

### S3 Supporting Information 3

#### S3.1 Theory-to-intervention tests

Deep-learning structure predictors such as AlphaFold have delivered a major breakthrough in recovering near-native, thermodynamically stable states. More recently, alternative-structure approaches have aimed to generate additional metastable conformations that may be functionally relevant. However, coverage of the metastable landscape remains incomplete, and without an accompanying mechanism it is difficult to determine which predicted alternatives matter and how exchange occurs.[10] The central frontier—and a prerequisite for functional interpretation—is therefore dynamics: whether an interconversion is kinetically accessible on relevant timescales, which pathway is taken through configuration space, and which microscopic elements control barrier crossing.

This work addresses that gap by shifting the emphasis from “which structures?” to “which transitions, which motions, and which players?”. Using the path-conditioned dimensionality-reduction framework introduced in Theory, high-dimensional dynamical data are converted into a transition-specific, testable mechanistic description: a low-dimensional representation of the barrier-crossing motion together with a ranked set of structural players that are most informative for that transition. Because informativeness alone does not establish causal leverage, the ranking is complemented by a separate intervention-tested causal analysis designed to identify which highly informative players also exhibit transition-specific dynamical leverage. This provides a physical basis for evaluating predicted alternative conformations beyond static accuracy metrics, and for formulating actionable hypotheses about how to predict, validate, and ultimately control the functional transition. Eight cohorts were defined according to the region of interest (ROI) used in REST2 scaling scheme and metadynamics (Table S3-1) for validation and intervention tests.

#### S3.2 Additional CV-based free-energy representations

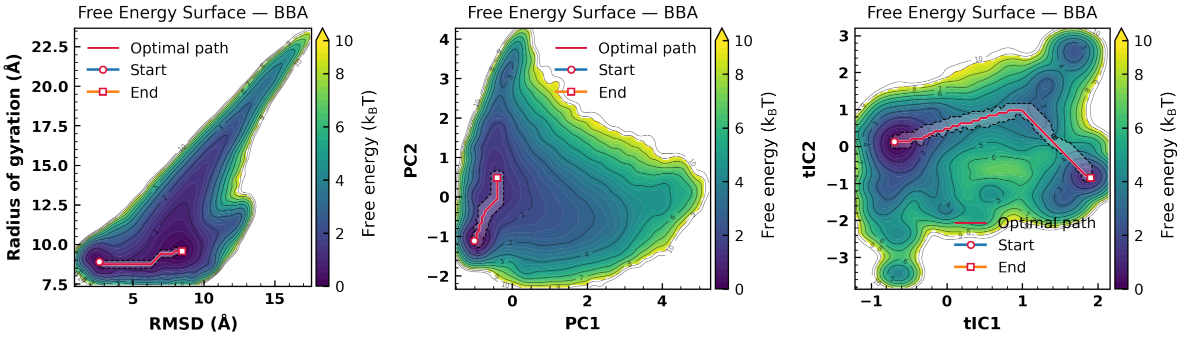

(a) BBA

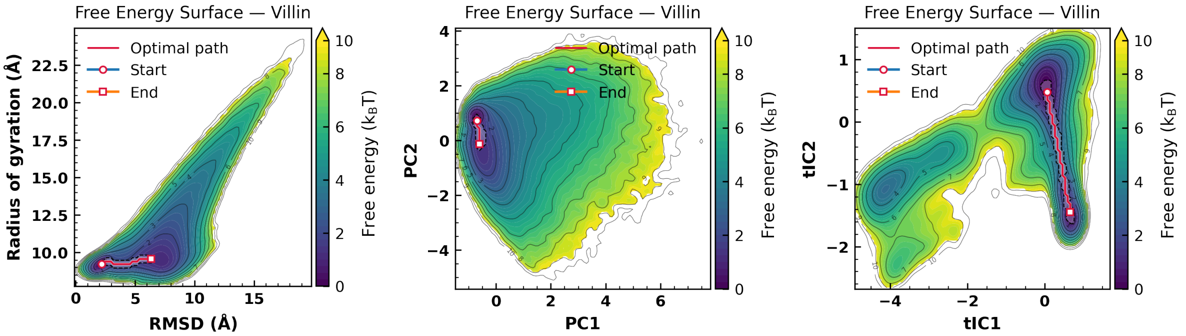

(b) Villin

Figure S3-1: Free energy diagram based on RMSD/RG, PCA and tICA for (a) BBA (325 microsecond), and (b) Villin (125 microsecond)

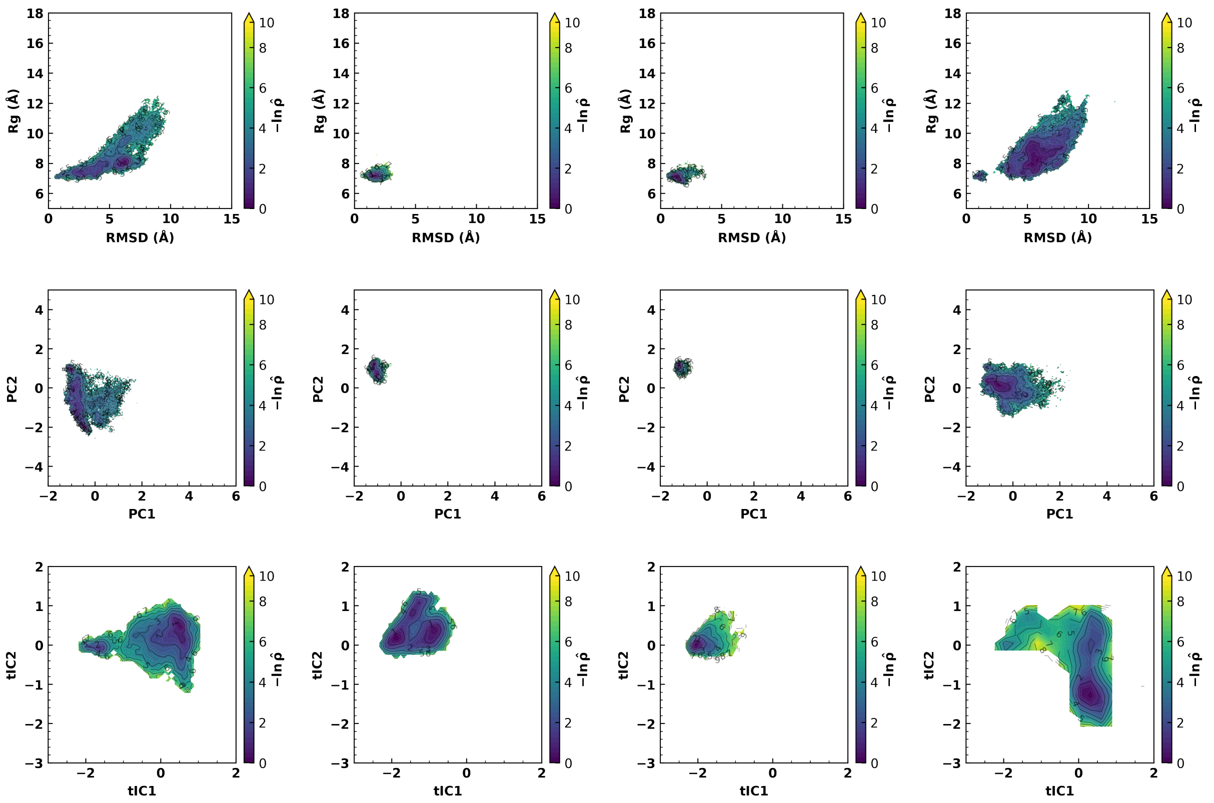

(a) (b) (c) (d)

Figure S3-2: RMSD/RG, PCA, and tICA based sampling-density maps of Trp-cage for (a) selected hotspots (Hot-ROI) (b) Random atoms (Random-ROI) (c) only PCA coefficient selected atoms (Path-PCA-ROI) (d) only tICA coefficient selected atoms (Path-tICA-ROI)

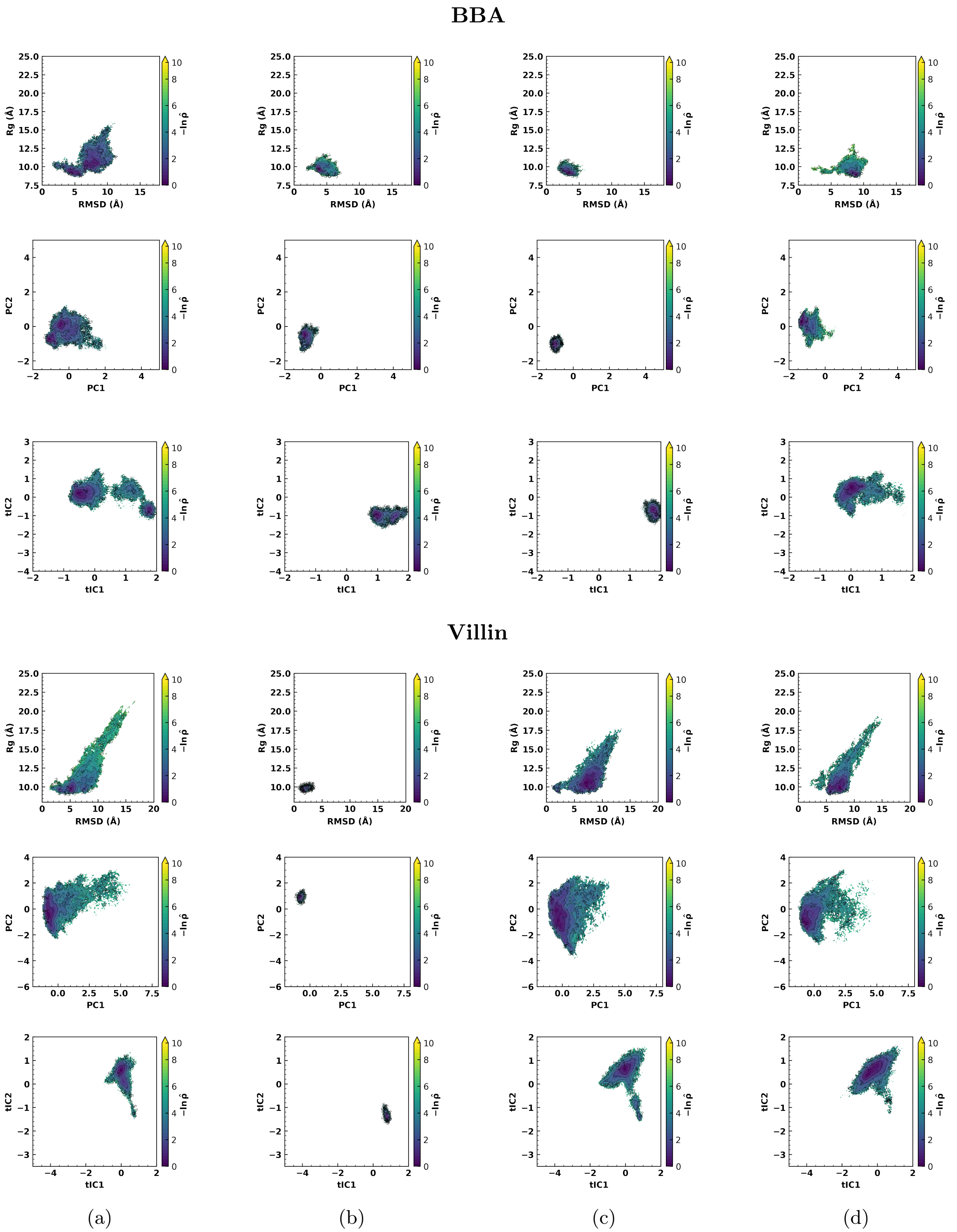

Figure S3-3: RMSD/RG, PCA, and tICA based sampling-density maps for (a) selected hotspots (Hot-ROI) (b) Random atoms (Random-ROI) (c) only PCA coefficient selected atoms (Path-PCA-ROI) (d) only tICA coefficient selected atoms (Path-tICA-ROI). Top block: BBA. Bottom block: Villin.

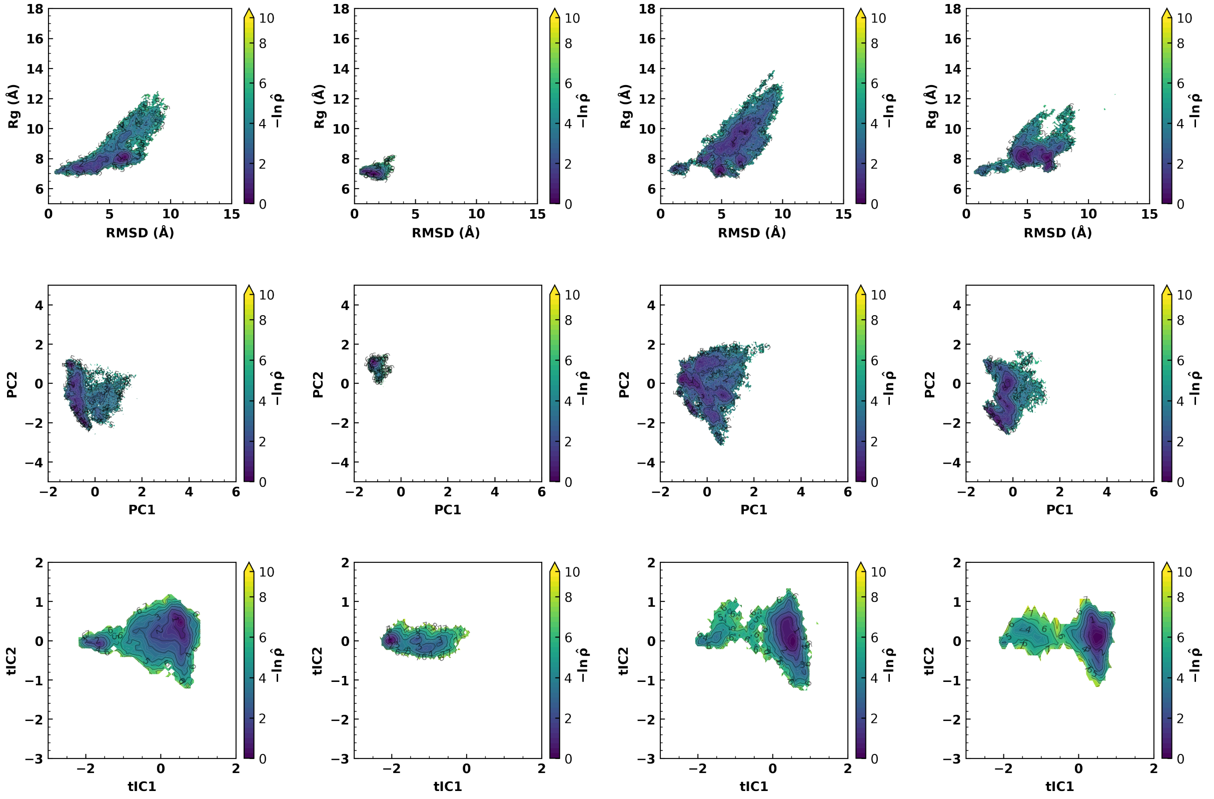

(a) (b) (c) (d)

Figure S3-4: RMSD/RG, PCA, and tICA based sampling-density maps of Trp-cage for (a) 8 percent selected hotspots (Hot-ROI) (b) 4 percent hotspots atoms (Hot-ROI) (c) only whole trajectory tICA coefficient selected atoms (G-tICA-ROI) (d) secondary selection hotspot (Tail-ROI)

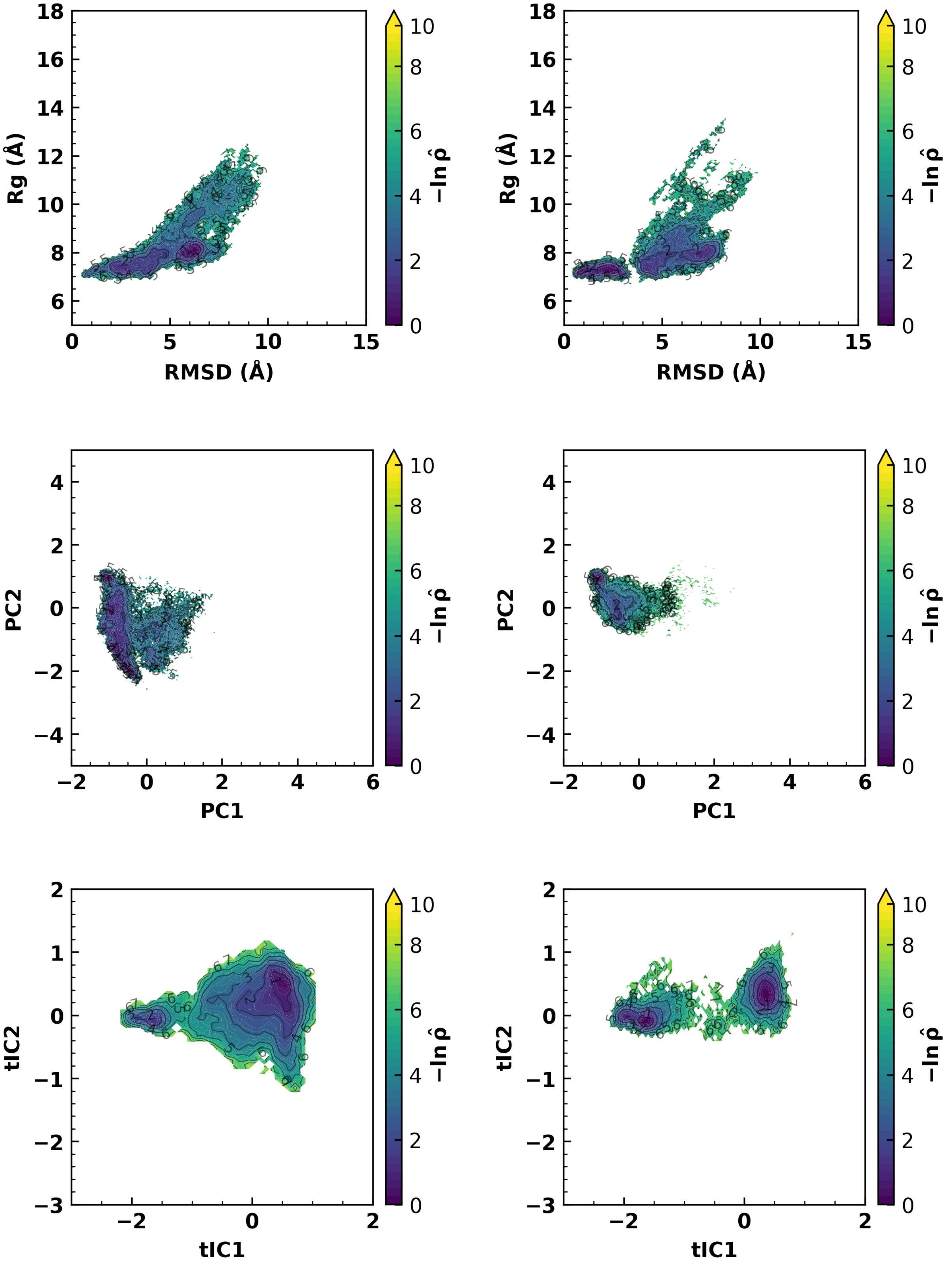

(a)

(b)

Figure S3-5: RMSD/RG, PCA, and tICA based sampling-density maps of Trp-cage for selected hotspots (Hot-ROI) based on (a) 100 percent simulation data (b) 20 percent simulation data

#### S3.3 Time-resolved transition trajectories in the reduced representation

To complement the sampling-density maps, the intervention trajectories were also examined in time-resolved form in the same reduced representation. This view makes it possible to distinguish repeated corridor-guided transitions from trajectories that remain trapped in a metastable basin or explore off-corridor regions. Consistent with the free-energy analysis, Hot-ROI produced more frequent basin-to-basin transitions, showed less long-lived trapping in the metastable states, and most notably remained concentrated on the prescribed corridor. By contrast, the baseline ROI choices spent more time trapped in the metastable basins and, when transitions occurred, showed weaker loyalty to the corridor.

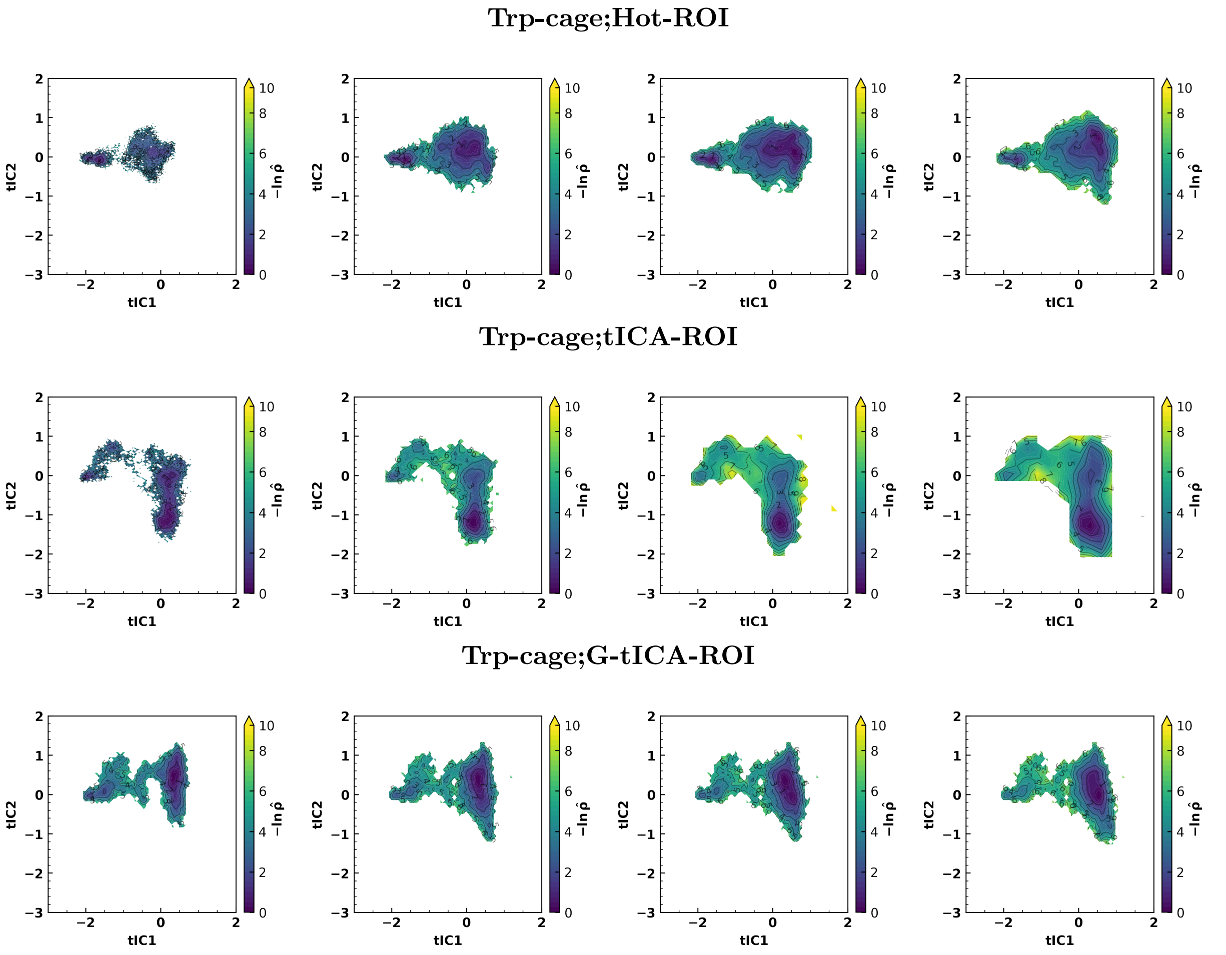

(a) (b) (c) (d)

Figure S3-6: tICA-based sampling-density maps of Trp-cage computed from (a) 20 ns (b) 40 ns (c) 60 ns (d) 100 ns simulations for selected hotspots (Hot-ROI), only tICA coefficient selected atoms (Path-tICA-ROI), and only whole trajectory tICA coefficient selected atoms (G-tICA-ROI).

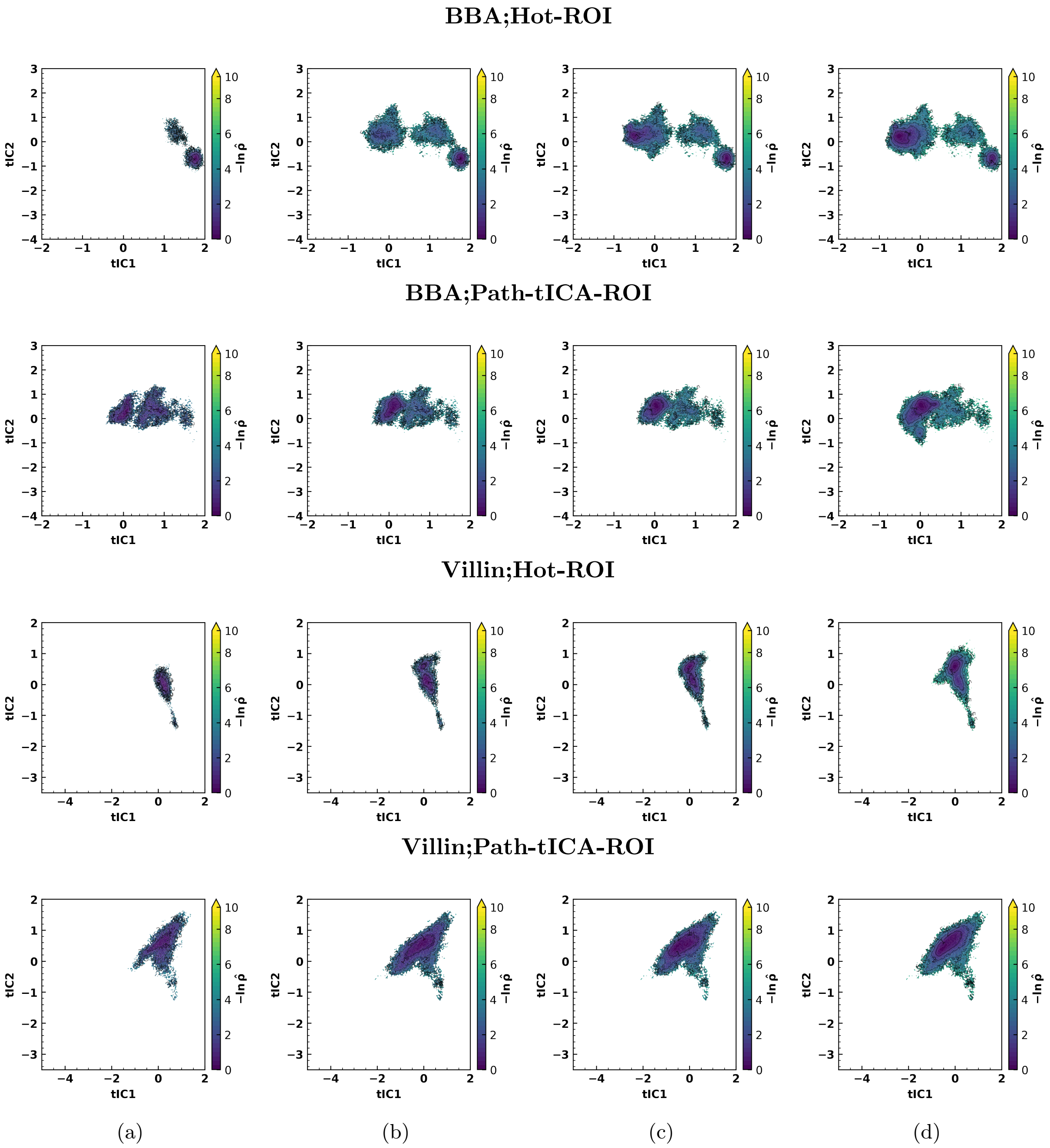

Figure S3-7: tICA-based sampling-density maps of BBA and Villin computed from (a) 20 ns (b) 40 ns (c) 60 ns (d) 100 ns simulations for selected hotspots (Hot-ROI), and only tICA coefficient selected atoms (Path-tICA-ROI).

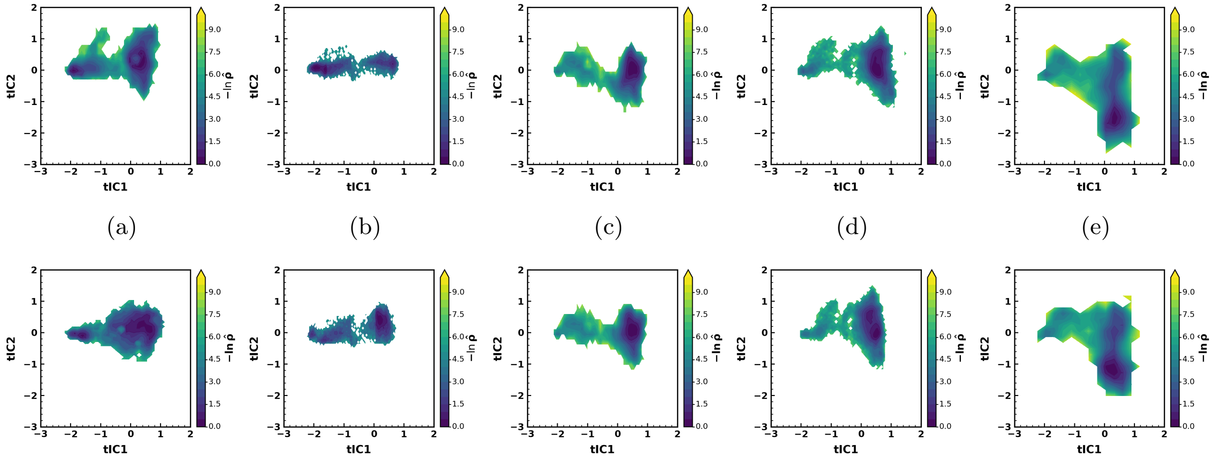

Figure S3-8: Additional independent realizations used in the quantitative corridor-fidelity analysis: (a) Hot-ROI, (b) 20% data-fraction, (c) Tail-ROI, (d) G-tICA-ROI, and (e) Path-tICA-ROI. For each condition, the upper and lower panels correspond to two different independent seeds.

#### S3.4 Early-time metadynamics sampling

To assess how rapidly each biasing choice expands sampling during metadynamics, RMSD–$R_g$ distributions were examined at early simulation time. All-backbone and Random-ROI biasing already produced broad displacements across the RMSD–$R_g$ plane, indicating rapid but weakly selective exploration. In contrast, Hot-ROI biasing expanded sampling in a more localized and transition-directed manner, consistent with acceleration that remains closer to the transition pathway.

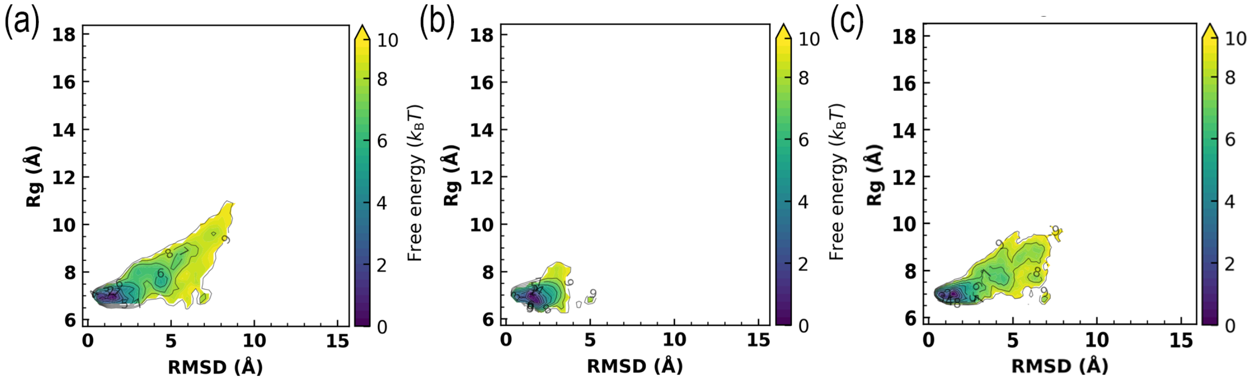

Figure S3-9: RMSD–$R_g$ free-energy profiles from metadynamics for early time simulation under three biasing choices: (a) all atoms, (b) Hot-ROI, and (c) Random-ROI.
